# Synchronous and opponent thermosensors use flexible cross-inhibition to orchestrate thermal homeostasis

**DOI:** 10.1101/2020.07.09.196428

**Authors:** Luis Hernandez-Nunez, Alicia Chen, Gonzalo Budelli, Vincent Richter, Anna Rist, Andreas S. Thum, Mason Klein, Paul Garrity, Aravinthan D.T. Samuel

**Affiliations:** Department of Physics, Harvard University, Cambridge, MA 02138, USA; Center for Brain Science, Harvard University, Cambridge, MA 02138, USA; Systems, Synthetic, and Quantitative Biology PhD program, Harvard University, Cambridge, United States; Harvard College, Harvard University, Cambridge, MA 02138, USA; National Center for Behavioral Genomics, Brandeis University, Waltham, United States; Department of Biology, Brandeis University, Waltham, United States; Volen Center for Complex Systems, Brandeis University, Waltham, United States; Department of Physics, University of Miami, Coral Gables,FL, United States; University of Leipzig, Institute of Biology, Talstraße 33, 04103 Leipzig, Germany

## Abstract

Body temperature homeostasis is an essential function that relies upon the integration of the outputs from multiple classes of cooling- and warming-responsive cells. The computations that integrate these diverse outputs to control body temperature are not understood. Here we discover a new set of Warming Cells (WCs), and show that the outputs of these WCs and previously described Cooling Cells (CCs^1^) are combined in a cross-inhibition computation to drive thermal homeostasis in larval *Drosophila*. We find that WCs and CCs are opponent sensors that operate in synchrony above, below, and near the homeostatic set-point, with WCs consistently activated by warming and inhibited by cooling, and CCs the converse. Molecularly, these opponent sensors rely on overlapping combinations of Ionotropic Receptors to detect temperature changes: Ir68a, Ir93a, and Ir25a for WCs; Ir21a, Ir93a, and Ir25a for CCs. Using a combination of optogenetics, sensory receptor mutants, and quantitative behavioral analysis, we find that the larva uses flexible cross-inhibition of WC and CC outputs to locate and stay near the homeostatic set-point. Balanced cross-inhibition near the set-point suppresses any directed movement along temperature gradients. Above the set-point, WCs mediate avoidance to warming while cross-inhibiting avoidance to cooling. Below the set-point, CCs mediate avoidance to cooling while cross-inhibiting avoidance to warming. Our results demonstrate how flexible cross-inhibition between warming and cooling pathways can orchestrate homeostatic thermoregulation.

---

Body temperature impacts all physiological processes and thus must be tightly controlled. In mammals, the preoptic area of the hypothalamus functions as a thermostat by combining the outputs of multiple warming- and cooling-activated cells to regulate physiological and behavioral mechanisms that keep body temperature near the homeostatic set-point.^2–4^ The computations that integrate warming and cooling cell outputs are not understood. The use of multiple physiological thermoregulatory mechanisms (e.g. evaporation of sweat for cooling and cutaneous vasoconstriction for warming) and behavioral mechanisms (navigation toward regions with warmer or colder ambient temperatures) makes it difficult to isolate the computations underlying thermoregulation in mammals.

Poikilotherms such as reptiles, fish, and insects lack physiological mechanisms to substantially warm or cool their bodies.^5^ These animals rely on behavior to locate regions with ambient temperatures closer to their homeostatic set-point. To do this, many poikilotherms, including cave beetles^6^ and larval *Drosophila*,^1^ have evolved exquisite neural and behavioral thermosensitivities (around 0.005°C/s), making them ideal to study the computations underlying thermal homeostasis.

The context in which cooling and warming must be interpreted in the brain is dependent on ambient temperature. When ambient temperature is below the homeostatic set-point, cooling should evoke avoidance behaviors; when ambient temperature is above the homeostatic set-point, warming should evoke avoidance behavior; and at the homeostatic set point any avoidance behavior should be inhibited. Thus, the outputs of thermosensory cells must be integrated flexibly to achieve thermoregulation.

Previous studies, in *Caenorhabditis elegans*,^7^ larval and adult *Drosophila melanogaster*,^5^ larval zebrafish^8^ and rodents^9^ have focused on the physiology of single thermosensory cell types and their contribution to thermoregulation in a specific ambient temperature context. Deriving a computation that incorporates the flexibility needed in different contexts requires determining how an animal integrates the contributions of both cooling- and warming-responsive cells across all ambient temperatures.

Here, we investigate the sensory cells and computations that control larval *Drosophila* body temperature. Previous work uncovered three cooling cells (CCs) in the Dorsal Organ Ganglion (DOG). The CCs are sensitive to temperature changes at ambient temperatures from 14°C to 34°C. The CCs are required for cooling avoidance from as low as 14°C towards 24°C (the homeostatic set-point); however, they are not required for warming avoidance above 24°C.^1^

Understanding homeostatic thermoregulation in larval *Drosophila* requires identifying the warming-responsive counterparts of the CCs and understanding how the outputs of cooling and warming cells are combined to make behavioral decisions at all ambient temperatures. Here, we uncover warming cells (WCs) with close morphological and genetic similarity to the CCs. Using optogenetics, calcium imaging, precise temperature control, sensory receptor mutants, and quantitative behavioral analysis, we derive a computation that uses ambient temperature context-dependent cross-inhibition between the simultaneous outputs of WCs and CCs. Flexible cross-inhibition allows the net effect of WC and CC outputs to drive cooling avoidance below 24°C, suppress avoidance to temperature changes near 24°C, and drive warming avoidance above 24°C. Our study reveals how simultaneously active opponent sensors are integrated in a context-dependent manner to achieve homeostatic regulation.

## Identifying warming cells

To identify the warming-responsive counterparts of the cooling cells (CCs), we used *in vivo* calcium imaging, expressing GCaMP6m^10^ under the control of the *pebbled-Gal4* driver that labels all anterior sensory cells in the larva^11^ (Fig. 1A). We subjected larvae to sinusoidal temperature waveforms, volumetrically imaged all anterior sensory ganglia, and used constrained non-negative matrix factorization (CNMF)^12^ to analyze activity patterns for evidence of temperature-sensitive cells (Fig. 1B). CNMF uncovered two novel warming cells (WCs) in the DOG, and identified the three previously described CCs (Fig. 1C). No other temperature-sensitive cells were apparent in any anterior sensory ganglia (Supplementary Fig. 1).

**Figure 1.**
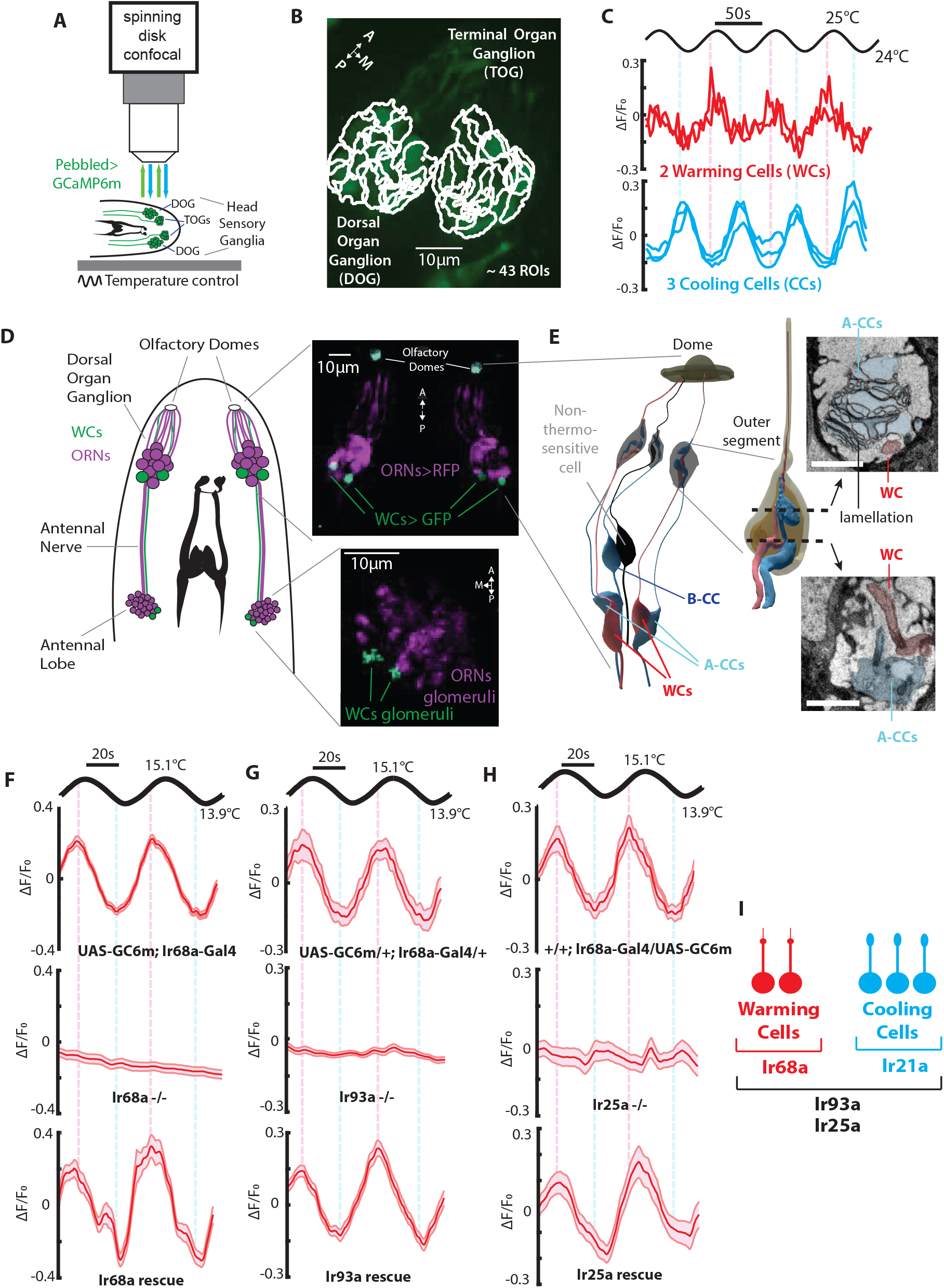
The cellular and molecular basis of warming sensing. **A**, Schematic representation of the larva head with all the anterior sensory organs expressing GCaMP6m via pebbled-Gal4 exposed to temperature sinusoidal fluctuations and imaged with a spinning disk confocal microscope.(Genoype: w^1118^;Pebbled-Gal4/UAS-GCaMP6m, n=6). **B**, CNMF segmentation of the regions of interest (ROIs) in the *Drosophila* larva’s head expressing GCaMP6m in the DOG and TOG. **C**, Responses recovered via CNMF of the CCs (in cyan) and the new WCs (in red). **D**,**E**, Anatomy of the WCs. **D**,Larvae expressing GFP in the WCs and RFP in the ORNs (UAS-GFP;Ir68a-Gal4/orco-RFP) imaged in the DOG (cell bodies) and the antennal lobe (axon terminals). **E**, Electron microscopy reconstruction of the thermosensory dendritic bulbs shared by CCs and WCs (scale bars= 1*μ* m). The top inset shows a section in the lamellated outer segment and the bottom inset shows an unlamellated part of WCs and CCs dendritic processes before the outer segment. **F-H**, Ir68a, Ir93a, and Ir25a are required for warming sensing. Fluorescence changes in the WCs of larvae with different genotypes exposed to a sine wave of temperature. **F**, Wildtype: UAS-GCaMP6m;Ir68a-Gal4 (n=8 animals), Ir68a defective mutants: UAS-GCaMP6m; Ir68a^PB^,Ir68a-Gal4 (n=8 animals), Ir68a rescue: UAS-GCaMP6m;(Ir68a^PB^,Ir68a-Gal4)/(Ir68a^PB^,UAS-Ir68a) (n=10 animals). **G**, Wildtype: UAS-GCaMP6m/+;Ir68a-Gal4/+ (n=8 animals), Ir93a defective mutants: UAS-GCaMP6m;Ir93a^MI05555^, Ir68a-Gal4/Ir93a^MI05555^ (n=14 animals), Ir93a rescue: UAS-CaMP6m/+;(Ir93a^MI05555^, Ir68a-Gal4)/(Ir93a^MI05555^, UAS-Ir93a) (n=8 animals). **H**, Wildtype: +;Ir68a-Gal4/UAS-GCaMP6m (n=8 animals), Ir25a defective mutants: Ir25a^2^;Ir68a-Gal4/UAS-GCaMP6m (n=20 animals), Ir25a rescue: (Ir25aBAC, Ir25a^2^)/Ir25a^2^;Ir68a-Gal4/UAS-GCaMP6m (n=20 animals). Shaded regions are the s.e.m. **I**, Warming and Cooling receptors and co-receptors summary.

Thermosensory cells in many animals have specialized morphologies that presumably enhance temperature detection.^13–15^ We used confocal and electron microscopy to reconstruct the anatomy of the WCs and CCs to better understand their structural specializations. Both WCs and CCs are located in the DOG, which mostly contains olfactory receptor neurons that project to different glomeruli in the antennal lobe.^16^ We sought cell-specific labels for the WCs. CCs in both adult and larval *Drosophila* express Ionotropic Receptors.^17–19^ We screened Ir genes known to be expressed in the DOG, and found that Ir68a-Gal4 exclusively labels the WCs (Supplementary Fig. 2, 3 and Extended Methods). Cell-specific labeling of the WCs using GFP revealed that each WC projects to a distinct warming glomerulus (Fig. 1D). The CCs project to a single cooling glomerulus.^1^ All thermosensory glomeruli are located posterior and dorsal to the olfactory glomeruli.

The anatomy and location of the WCs and CCs facilitated their reconstruction using electron microscopy (Extended Methods). Consistent with previous studies,^1^ we refer to the posterior CCs as A-CCs, and to the most anterior CC as B-CC. The cell bodies and outer segments of the A-CCs and WCs are adjacent (Fig. 1E). The outer segments of the CCs and WCs have specialized morphologies, presumably containing signal transduction machinery. The CC outer segments are large and lamellated with heavily infolded plasma membranes. The WC outer segments are smaller and unlamellated (Fig. 1E inset). These anatomical features are consistent with those of the WCs and CCs of adult *Drosophila*.^15, 16^ The larval WCs, but not the CCs, also have a thin dendrite that protrudes to the surface of the olfactory dome (Fig. 1E, Supplementary Fig. 4, and Supplementary Material). The cell body and outer segment of the B-CC are adjacent to a non-thermosensitive cell of the DOG (Supplementary Fig. 1 and Supplementary Material).

## The molecular basis of warming sensing

Because *Ir68a-Gal4* labels the WCs, we asked whether *Ir68a* might directly contribute to their thermosensitivity. Consistent with *Ir68a* expression in the WCs, a Gal4 reporter under the control of the endogenous *Ir68a* promoter (*Ir68a^T2A-Gal4^*) drove cell-specific expression in the WCs. *Ir68a* was required for WC responsiveness to warming, as a loss-of-function mutation in *Ir68a* (*Ir68a^PB^*) abolished temperature-evoked calcium dynamics in the WCs (Fig. 1F). The defect was specific, as cell-specific expression of wildtype *Ir68a* restored WC thermosensitivity in the *Ir68a^PB^* mutant (Fig. 1F).

Most Ionotropic Receptors in *Drosophila* appear to function as heteromers.^20, 21^ For example, the CCs require a set of three Ionotropic Receptors to respond to temperature changes: Ir21a, Ir93a, and Ir25a.^17, 18^ We used Ir21a^T2A-Gal4^ lines,^22^ in which Gal4 is expressed under the control of the endogenous Ir21a promoter, to drive GFP expression and found that Ir21a expression is specific to the CCs and was not detected in the WCs or elsewhere in the larva. For *Ir93a*, immunostaining revealed that the expression pattern of the Ir93a receptor is specific to the WCs and CCs (Supplementary Fig. 2). Ir25a is expressed in many anterior sensory cells including the WCs and CCs.^17, 23–25^ Consistent with their expression in WCs, mutations disrupting either of these two receptors, Ir93a (*Ir93a^MI^*) or Ir25a (*Ir25a^2^*), abolished WC thermosensitivity (Fig. 1G, H). The thermosensitivity of the WCs was restored in each mutant by cell-specific re-expression of the corresponding wildtype receptor (Fig. 1G, H).

Our results suggest a model in which distinct but overlapping sets of Ionotropic Receptors confer thermosensitivity to the WCs and CCs (Fig. 1I). Ir68a is specifically needed by the WCs to sense warming. Ir21a is specifically needed by the CCs to sense cooling. Ir93a and Ir25a are needed by both WCs and CCs to sense any temperature change. We further tested this model by ectopic expression of Ir68a and Ir21a in the CCs and WCs, respectively. Ectopic expression of Ir68a in the CCs diminished their sensitivity to cooling, while ectopic expression of Ir21a in the WCs transformed them into cooling sensors (Supplementary Fig. 5). These findings support a model where Ir68a and Ir21a generate opposite thermosensitive polarities in WCs and CCs.

## WCs and CCs are synchronous and opponent thermosensors

A first step to examine how WCs and CCs may be integrated to mediate homeostatic temperature control is to quantify the dynamics of their temperature evoked neural responses. We labeled WCs and CCs with GCaMP6m and measured their calcium responses to warming or cooling step stimuli. A warming step evoked a transient increase in WC calcium levels and a transient decrease in CC calcium levels (Fig. 2A). A cooling step evoked a transient decrease in WC calcium levels and a transient increase in CC calcium levels (Fig. 2B). The average time for the calcium response peak (*τ*_peak_) was not significantly different in WCs and CCs during warming or cooling steps (Fig. 2C). The time for adaptation of the neural response to baseline calcium levels (*τ*_adaptation_) was also not significantly different regardless of the polarity of the step stimulus (Fig. 2D). Thus, WCs and CCs are synchronous and bidirectional phasic sensors of temperature change with opposite polarity. WCs are activated by warming and inhibited by cooling, while CCs are activated by cooling and inhibited by warming (Fig. 2E).

**Figure 2.**
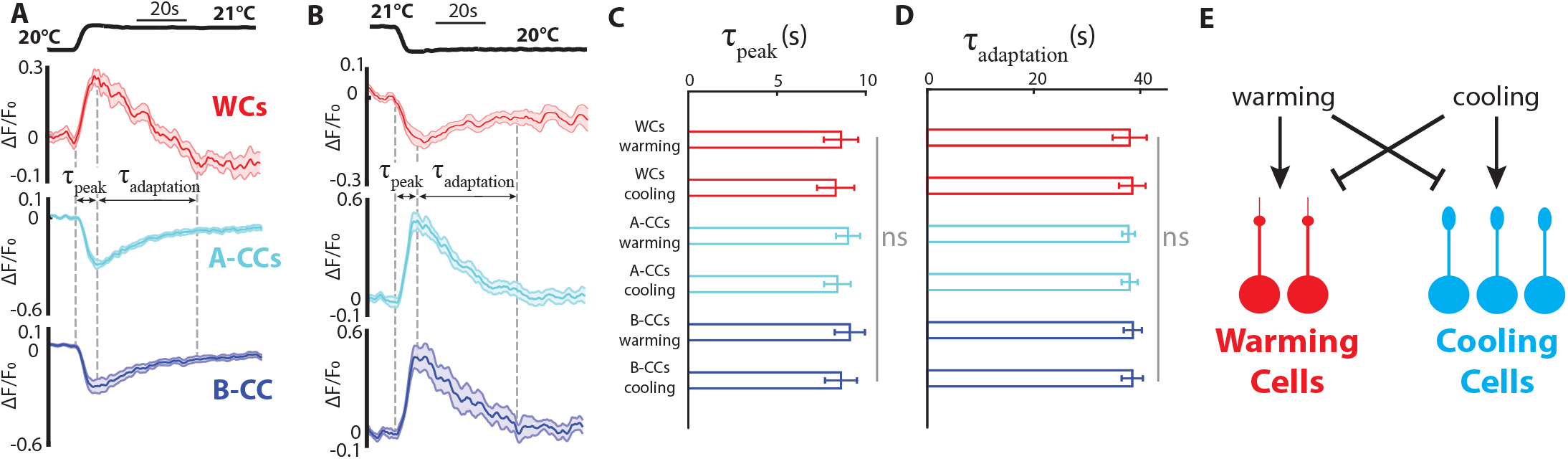
Synchronous and opponent sensors of temperature change. **A**, **B**, *Drosophila* larvae expressing GCaMP6m in the WCs (UAS-GCaMP6m;Ir68a-Gal4) or CCs (R11F02-Gal4; UAS-GCaMP6m) calcium responses to a 1°C temperature increase (**A**) or decrease (**B**). (WCs responses in red with n=12 larvae, and CCs responses in cyan (A-CCs) and blue (B-CCs) with n=10, the shaded regions are the s.e.m.). **C**, The peak times of WCs and CCs responses to the warming and cooling step stimuli are not significantly different (Kruskal-Wallis test, error bars are the s.e.m). **D**, The adaptation times of WCs and CCs responses to the warming and cooling step stimuli are not significantly different (Kruskal-Wallis test, error bars are the s.e.m). The grey ‘ns’ in **C** and **D** indicates no statistically significant difference. **E**, WCs are activated by warming and inhibited by cooling. CCs are inhibited by warming and activated by cooling.

The synchronous and opponent calcium responses of the WCs and CCs are not specific to step stimuli. The WCs and CCs also display similar response properties when stimulated with sinusoidal temperature fluctuations (Supplementary Fig. 6).

## Behavioral flexibility is not encoded in WCs or CCs neural responses

The homeostatic temperature set-point of first and second instar larval *Drosophila* is near 24°C. To achieve thermoregulation, larvae must flexibly interpret warming and cooling in different contexts of ambient temperature (below, near, or above 24°C). Understanding the origin of this flexibility requires mapping the interplay between ambient temperature, temperature change, thermosensory neuron activity and behavioral responses at different ambient temperatures. To do this, we developed a temperature control technique to deliver the same temperature waveforms at multiple ambient temperatures during calcium imaging and behavioral experiments (Fig. 3A, B, Supplementary Fig. 7 and Extended Methods). We used this setup to quantify the temperature-evoked responses of freely behaving larvae. Larval *Drosophila* navigate by alternating periods of forward crawling called runs and reorientation events called turns^26^ (Fig. 3C). Using an unsupervised classifier to segment behavioral sequences (Supplementary Fig. 8, and Extended Methods) we verified that turning rate (the probability of turning per time unit) is the only motor program significantly modulated by temperature changes. Turning is the principal avoidance response that helps larvae find more favorable orientations, capturing the valence, dynamics, and intensity of the behavioral response to a temperature stimulus.^1, 26, 27^

**Figure 3.**
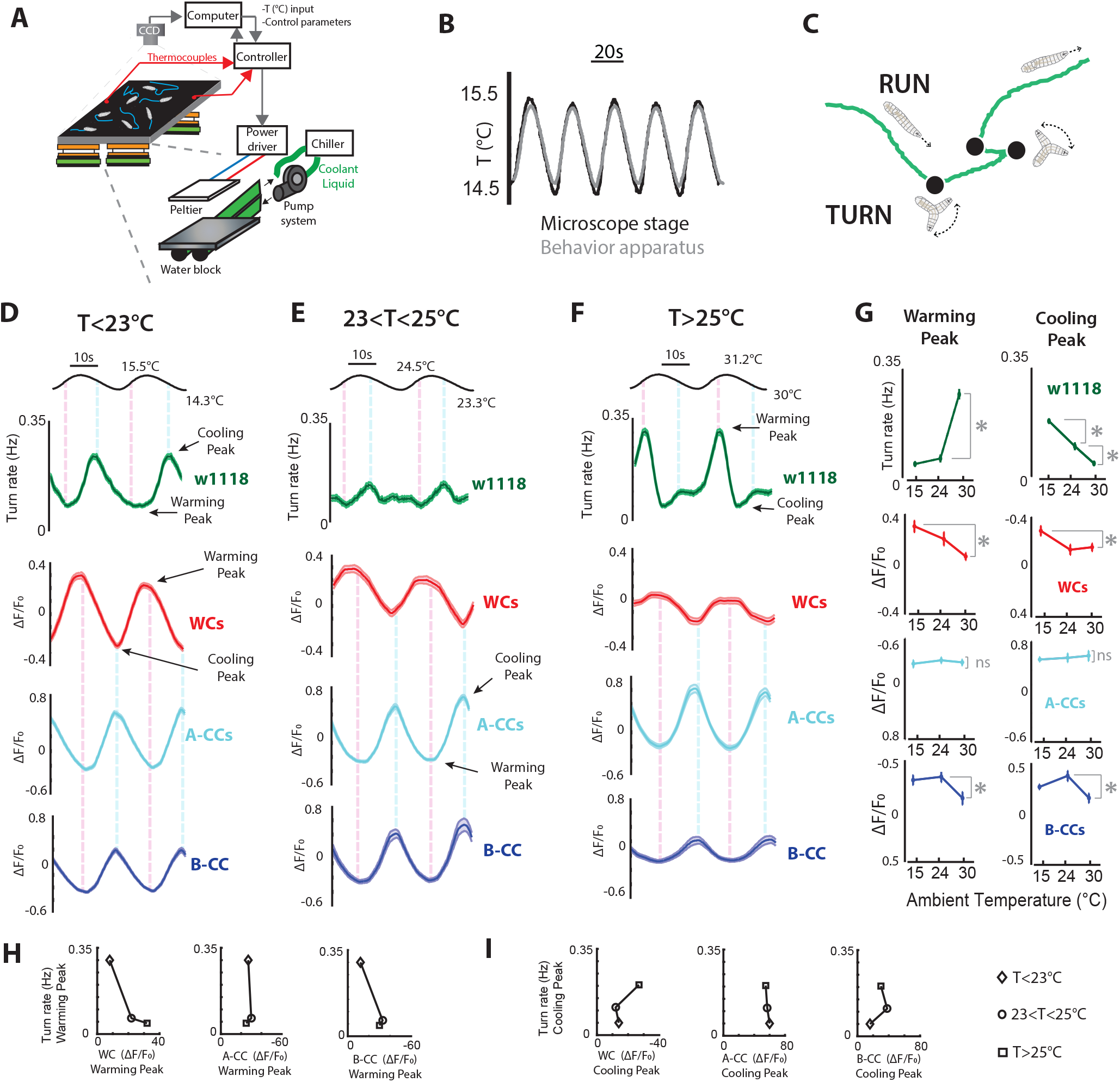
Behavioral flexibility is not encoded in WCs or CCs neural responses. **A**, Temperature control apparatus to study larval behavior. Four thermoelectric Peltier elements heat or cool a copper plate, rectangular agar is deposited on the copper plate and larva crawl on the agar. Water blocks are attached to the thermoelectric elements and flow antifreeze at 5°C to cool the thermoelectric elements. Thermocouples are used to measure temperature on the agar surface. A high pixel density CCD camera with an infrared filter is used to record larval movements and four sets of infrared LED bars provide dark field illumination from the sides. **B**, Sinusoidal waves of temperature measured on the top of the agar of the behavior rig (in grey) and on top of the microscope stage where we conduct calcium imaging (in black). **C**, Larvae navigate by alternating periods of forward crawling called runs (green lines) and reorientation events called turns (Black dots). **D-F**, Behavioral and neural responses to sinusoidal temperature fluctuations below (**D**), near (**E**), and above (**F**) the homeostatic set-point. In the first row are the experimental results of the turning rate responses to sinusoidal waves of temperature of wildtype larvae (w^1118^ in green, shaded regions are the s.e.m.). In the second row are the fluorescence changes in the WCs of Ir68a-Gal4; UAS-GCaMP6m larvae exposed to sinusoidal waves of temperature (red curves, n=8-10 animals, shaded regions are the s.e.m.). In the third row are the fluorescence changes in the A-CCs of R11F02-Gal4; UAS-GCaMP6m larvae exposed to sinusoidal waves of temperature (cyan curves, n=7-10 animals, shaded regions are the s.e.m.). In the fourth row are the fluorescence changes in the B-CCs of R11F02-Gal4; UAS-GCaMP6m larvae exposed to sinusoidal waves of temperature (blue curves, n=7-10 animals, shaded regions are the s.e.m.).**G**, Summary of the peak behavioral and neural responses to cooling and warming. (First row) The turning rate response to warming is larger at high ambient temperatures and neutral at low temperatures and at the homeostatic set-point. The turning rate response to cooling is larger at low temperatures, and lower at high temperatures. (Second row) The WCs’ positive peak response to warming and negative peak response to cooling are larger below and near the homeostatic set-point and closer to zero at high temperatures. (Third row) The A-CCs positive peak response to cooling and negative peak response to warming are not significantly different at all temperature baselines. (Fourth row) The B-CCs’ positive peak response to cooling and negative peak response to warming are closer to zero at high temperatures. The error bars are s.e.m. of the individual behavioral or neural responses averaged to calculate the responses displayed in **D**,**E**, **F**. The * indicate that the peak responses between the brackets are different with Kruskal-Wallis test p<0.01. The grey ‘ns’ indicates no statistically significant difference. **H**, **I**, The WCs’ and CCs’ peak responses to warming (**H**) and cooling (**I**) versus the peak behavioral responses at all ambient temperatures.

To analyze how cooling and warming modulate turning rate at different ambient temperatures, freely moving larvae were exposed to identical sinusoidal waves of 1.2°C amplitude centered at temperatures below (14.9°C), near (23.9°C), or above (30.6°C) the homeostatic set-point. Below the set-point, turning rate peaks during the cooling phase (Fig. 3D), consistent with cooling avoidance behavior. Near the set-point, avoidance responses are suppressed (Fig. 3E). Above the set-point, turning rate peaks during warming phase (Fig. 3F), consistent with warming avoidance behavior. Thus, warming and cooling are interpreted differently at different ambient temperatures.

We asked whether the flexible computation that transforms warming and cooling into behavior reflects differences in the sensitivity of WCs and CCs at different ambient temperatures. We tested the WCs and CCs neural responses to the same sinusoidal waves used to measure behavioral responses. Near the homeostatic set-point, where avoidance behavioral responses are suppressed, WCs are activated during the warming phase and inhibited during the cooling phase, whereas the CCs are activated during the cooling phase and inhibited during the warming phase (Fig. 3E). Although neither temperature change should evoke avoidance near 24°C, WCs and CCs still display strong, opponent physiological responses to temperature change in this range. At all ambient temperatures, the CCs are activated by cooling and inhibited by warming with similar sensitivities (Fig. 3D-G), only with a sensitivity decrease of the B-CCs at high temperatures (Fig. 3D-F). Thus, the CCs do not appear to become more sensitive at low temperatures to upregulate cooling avoidance (Fig. 3I). Similarly, the WCs are activated by warming and inhibited by cooling at all temperatures (Fig. 3D-G), and are even more sensitive at low temperatures (Fig. 3G). Likewise, the WCs do not appear to become more sensitive at high temperatures to upregulate warming avoidance (Fig. 3H). Therefore, the flexibility in the computation underlying thermal homeostasis is not encoded in the physiological thermosensitivity of WCs and CCs.

## Candidate computations underlying thermal homeostasis

How are the WC and CC outputs combined in a computation for thermal homeostasis? Answering this question requires measurement of the transformation from WC and CC activity into changes in turning rate. In linear systems, reverse-correlation analysis can reveal the filters that transform neural activity into behavior.^27^ To perform reverse-correlation analysis, we recorded freely moving larvae responding to optogenetic white-noise stimulation of either their WCs or CCs (Fig. 4A, and Extended Methods). The filters that transform WC and CC activity into turning rate were identical (Fig. 4B), suggesting that functionally similar pathways transform the activities of WCs and CCs into behavior.

**Figure 4.**
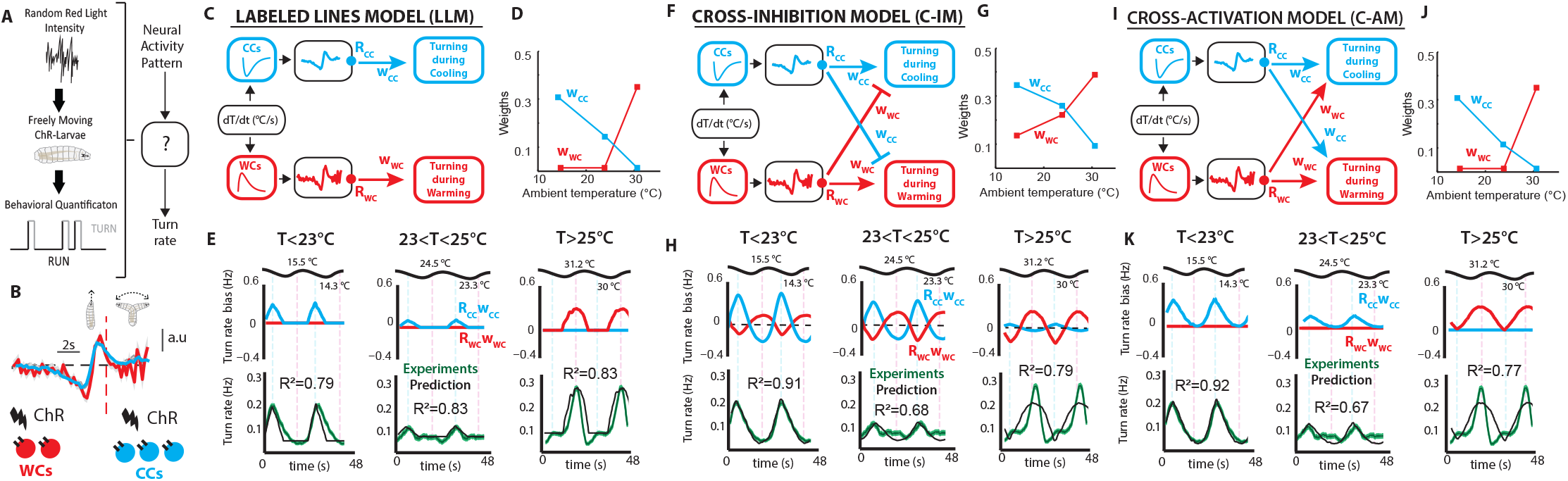
Candidate computations underlying thermal homeostasis. **A**, Mapping sensory neuron activity patterns to behavior. The mathematical function that converts an optogenetic stimulus pattern in a specific type of sensory cells and behavior can be estimated using reverse-correlation of behavioral responses to white noise optogenetic stimulation. In our case this is the transformation from optogenetic activation to turning rate. **B**, Linear filters obtained with white noise optogenetic stimulation of WCs (curve in red) and CCs (curve in cyan). We abbreviated CsChrimson as ChR. **C**, **F**, **I**, Schematic representation of the candidate computations underlying thermal homeostasis. The WC and CC outputs are convolved with behavior filters (Extended Methods) and the filter outputs (R_CC_ and R_WC_) are then combined linearly with the scalar weights w_CC_ and w_WC_. There are three possible symmetric architectures that linearly combine the outputs of WCs and CCs: (i) The *Labeled Lines Model (LLM)*, where turning during cooling is exclusively regulated by the CCs and turning during warming is exclusively regulated by the WCs (**C**); (ii) the *Cross-Inhibition Model (C-IM)*, where WCs can cross-inhibit turning during cooling in addition to driving turning during warming, and CCs can cross-inhibit turning during warming in addition to driving turning during cooling (**F**); (iii) the *Cross-Activation Model (C-AM)*, where both neuron types can drive both turning during cooling and warming (**I**). **D**, **G**, **J**, We used linear-regression to obtain the weights w_CC_ and w_WC_ of the contributions of CCs and WCs to turning rate in each model architecture. **E**, **H**, **K**, (First row) The outputs of the behavioral filter of the warming pathway (*R_WC_*) and the cooling pathway (*R_CC_*), scaled by their respective weights (*w_WC_* and *w_CC_*). This shows the contribution of the WCs and CCs during cooling and warming for each model architecture. (Second row) Experimental results of the turning rate response to sine waves of temperature of wildtype larvae (w^1118^ in green) and model predictions (in black) for each architecture. Shaded regions around the green curve are then s.e.m. Correlations coeficients between the model predictions and experimental results are reported in each curve.

Any candidate linear model for combining WC and CC activity to produce behavior is fully determined by knowing how sensory inputs map to WC and CC activity, how WC and CC activities separately map to turning rates, and how sensory inputs map to turning rate. Our datasets include all of these measurements. The remaining task is to determine the architecture of the model and the scalar weights (*w_WC_* and *w_CC_*) that determine the magnitude of the contributions of WCs and CCs to turning during warming or cooling.

In principle, there are only three symmetric model architectures posited to integrate WC and CC outputs. In the *Labeled Lines Model* (*LLM*), the WCs exclusively drive behavior during warming, and the CCs exclusively drive behavior during cooling (Fig. 4C). Alternatively, the WCs might cross-modulate behavior during cooling or the CCs might cross-modulate behavior during warming. The polarity of the cross-modulation can be either negative in a *Cross-Inhibition Model* (*C-IM*) (Fig. 4F), or positive in a *Cross-Activation Model* (*C-AM*) (Fig. 4I).

In each of the three alternative architectures, the only free variables are the weights of the outputs of the WCs (*w_WC_*) and the CCs (*w_CC_*) at different ambient temperatures. For each architecture, we used linear regression analysis to predict the necessary weights to account for the behavior of wild-type larvae. In the *LLM* (Fig. 4C), below and near the homeostatic set-point, *w_WC_* must be zero to prevent an increase in turning during warming, while *w_CC_* must scale the magnitude of turning during cooling (Fig. 4D, E left and center). Above the set-point, *w_WC_* scales the magnitude of turning during warming and *w_CC_* must be zero to prevent an increase in turning during cooling (Fig. 4D, E right). The *LLM* can accurately predict behaviors in all contexts (*R*^2^ > 0.79).

In the *C-IM* (Fig. 4F), below the homeostatic set-point, CC output is weighted more strongly than WC output to allow a net increase in turning during cooling (Fig. 4G, H left). Near the set-point, *w_WC_* and *w_CC_* values are not zero (like in the *LLM*), but are balanced so that the mutual inhibition of both pathways results in attenuated turning rates (Fig. 4G, H center). Above the set-point, WC output is weighted more strongly than CC output to allow a net increase in turning during warming (Fig. 4G, H right). The *C-IM* can also accurately predict behaviors in all contexts (*R*^2^ > 0.68).

In the *C-AM* (Fig. 4I), both WC and CC outputs regulate turning rate positively. As in the *LLM*, *w_WC_* has to be zero below and near the homeostatic-set-point to prevent turning increases during warming, while *w_CC_* has to scale the magnitude of turning during cooling (Fig. 4J, K left and center). Above the set-point, *w_WC_* scales the magnitude of turning during warming and *w_CC_* has to be zero to prevent an increase in turning during cooling (Fig. 4J, K right). The *C-AM* can also accurately predict behaviors in all contexts (*R*^2^ > 0.67).

All candidate models are capable of predicting wildtype larval *Drosophila* behavior at all ambient temperatures, albeit with different values of the weights of the WC and CC outputs. In each model, behavioral flexibility is encoded in the values of the weights (*w_WC_* and *w_CC_*) at different ambient temperatures. Determining which model represents the actual computation underlying thermal homeostasis requires independent manipulation of the neural activity of WCs and CCs at all ambient temperatures.

## Flexible cross-inhibition underlies thermal homeostasis

Thermosensory stimuli simultaneously affect both WC and CC activity, confounding attempts to distinguish the behavioral consequences of WC versus CC activation or inhibition in different ambient temperature contexts. To independently manipulate WC and CC activity, we turned to optogenetics. With cell-specific expression of CsChrimson^28^ in either WCs or CCs using controlled optogenetic illumination, we induced fictive temperature changes onto each cell type at different ambient temperature contexts and compared model predictions with experimental results.

We found that increasing optogenetic stimulation of WCs (fictive warming) caused an increase in turning rate at all ambient temperatures (Fig. 5A). This result supports the Cross-Inhibition Model (C-IM), where activation of WCs causes increases in turning rates at all temperatures. In contrast, the Labeled Lines Model (LLM) and the Cross-Activation Model (C-AM) fail to predict the turning rate increase caused by WCs activation below and near the homeostatic set-point (Fig. 5A). Decreasing optogenetic stimulation of WCs (fictive cooling), caused a decrease in turning rate at all ambient temperatures (Fig. 5B). This also supports the C-IM, where a decrease in WC activity inhibits turning rates at all temperatures. The LLM fails to predict turning rates at all temperatures, and the C-AM fails to predict turning rates below and near the homeostatic set-point (Fig. 5B).

**Figure 5.**
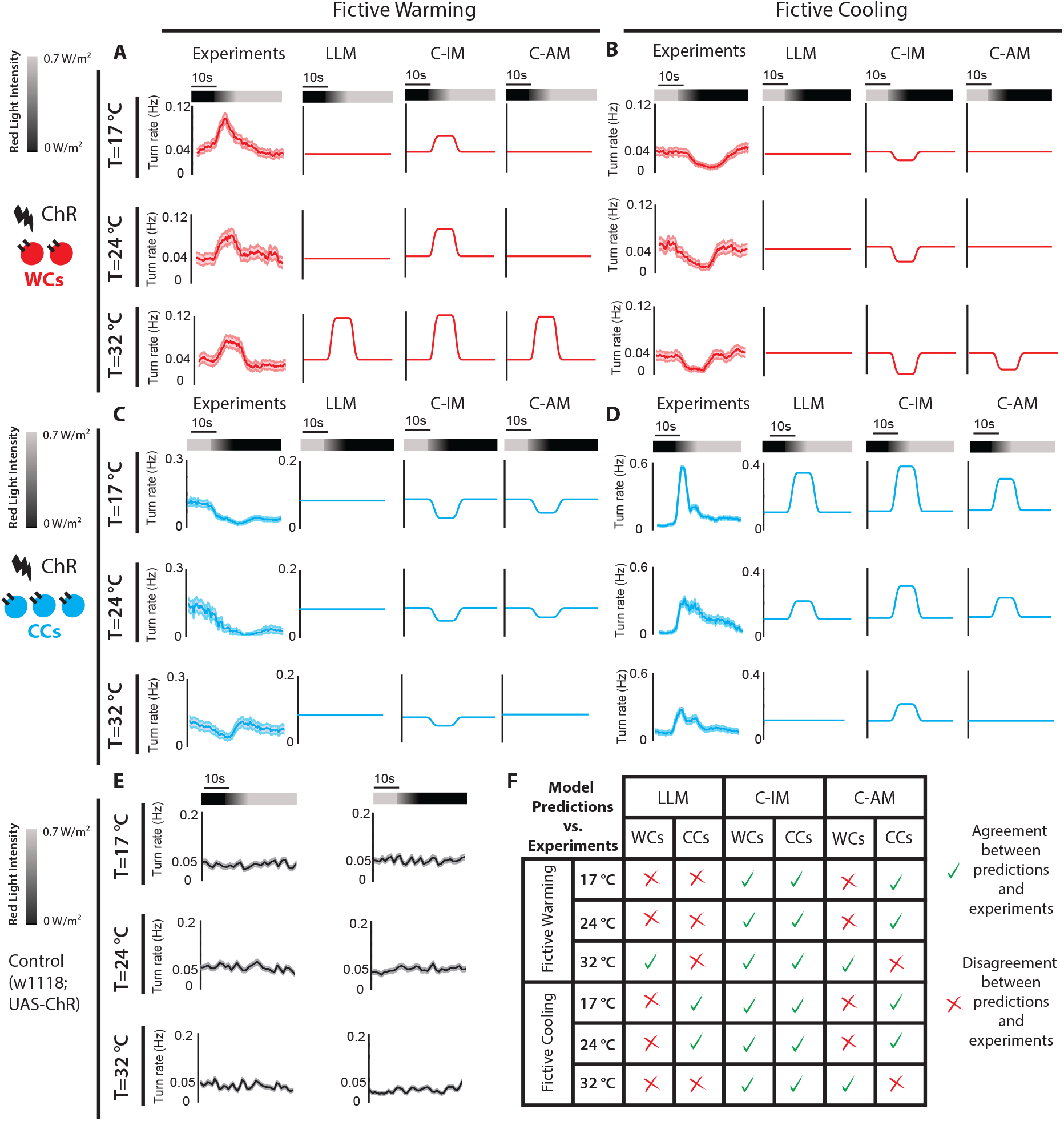
Flexible cross-inhibition underlies thermal homeostasis. **A**, Optogenetic fictive warming of WCs at different ambient temperatures. (First column) Turning rate of larvae expressing the red shifted optogenetic channel CsChrimson (ChR) in the WCs (w^1118^;UAS-CsChrimson/+;Ir68a-Gal4/+) exposed to positive ramps of red light (fictive warming) at all ambient temperatures. (Other columns) Model-predicted turning rates in response to a ramp increase of WCs’ activity. **B**, Optogenetic fictive cooling of WCs at different ambient temperatures. (First column) Turning rate of larvae expressing the red shifted optogenetic channel CsChrimson (ChR) in the WCs (w^1118^;UAS-CsChrimson/+;Ir68a-Gal4/+) exposed to negative ramps of red light (fictive cooling) at all ambient temperatures. (Other columns) Model-predicted turning rates in response to a ramp decrease of the WCs’ activity. In **A** and **B** n=116-125 animals for each ambient temperature and shaded regions are the s.e.m. **C** Optogenetic fictive warming of CCs at different ambient temperatures. (First column) Turning rate of larvae expressing the red shifted optogenetic channel CsChrimson (ChR) in the CCs (w^1118^;UAS-CsChrimson/+;R11F02-Gal4/+) exposed to negative ramps of red light (fictive warming) at all ambient temperatures. (Other columns) Model-predicted turning rates in response to a ramp decrease of the CCs’ activity. **D**, Optogenetic fictive cooling of CCs at different ambient temperatures. (First column) Turning rate of larvae expressing the red shifted optogenetic channel CsChrimson (ChR) in the CCs (w^1118^;UAS-CsChrimson/+;R11F02-Gal4/+) exposed to positive ramps of red light (fictive cooling) at all ambient temperatures. (Other columns) Model-predicted turning rates in response to a ramp increase of the CCs’ activity. In **C** and **D** n=90-101 animals for each ambient temperature and shaded regions are the s.e.m. **E**,Turning rate of control larvae (w^1118^; UAS-CsChrimson/+) fed with all-trans-retinal (ATR) and exposed to positive and negative ramps of red light. n=85-120 animals at each ambient temperature, shaded regions are the s.e.m. **F**, Summary chart of the agreement between model predictions and experimental results. Only the C-IM predictions agree with experimental results at all ambient temperatures.

We observed an opposite pattern with the CCs. Decreasing optogenetic stimulation of the CCs (fictive warming) caused a decrease in turning rate at all temperatures(Fig. 5C). This supports the C-IM, where a decrease in CC activity inhibits turning rate at all temperatures. The LLM fails to predict turning rates at all temperatures, and the C-AM fails to predict turning rates above the homeostatic set-point (Fig. 5C). Finally, increasing optogenetic stimulation of CCs (fictive cooling) caused an increase in turning rate (Fig. 5D). This also supports the C-IM model that predicts that CC activity increases are reflected in turning rate increases at all temperatures. The LLM and the C-AM fail to predict turning rates near or above the homeostatic set-point.

Control animals exhibited no behavioral responses to changes in optogenetic stimulation (Fig. 5E). The Cross-Inhibition Model is the only linear model that captures the contribution of WCs and CCs to thermoregulation in all contexts (Fig. 5F). This cross-inhibtion does not operate with fixed weights because the flexibility needed to interpret warming and cooling in different ambient temperature contexts is not encoded in the neural responses of WCs and CCs. Instead, the flexibility of the cross-inhibition computation is encoded in the changing values of the weights (*w_WC_* and *w_CC_*) that appropriately scale the contribution of warming and cooling pathways.

## Cross-inhibition computations in mutant larvae

The cross-inhibition computation explains how WC and CC outputs are effectively integrated to regulate behavior. We asked whether this computation can also explain the deficits in thermoregulation caused by non-functional WCs and/or non-functional CCs. Introducing non-functional mutations for receptors that are required for warming (*Ir68a^PB^*) or cooling sensing (*Ir21a^123^*) is equivalent to calculating the output of the cross-inhibition computation without WCs or CCs (Fig. 6A). Using a mutant for Ir93a (*Ir93a^MI^*) –a receptor only expressed in WCs and CCs and required for their thermosensitivity– is equivalent to calculating the output of the cross-inhibition computation in the absence of both WCs and CCs. Therefore, the experimentally measured behavioral responses of *Ir93a^MI^* mutants represent the baseline turning rate of the cross-inhibition computation.

**Figure 6.**
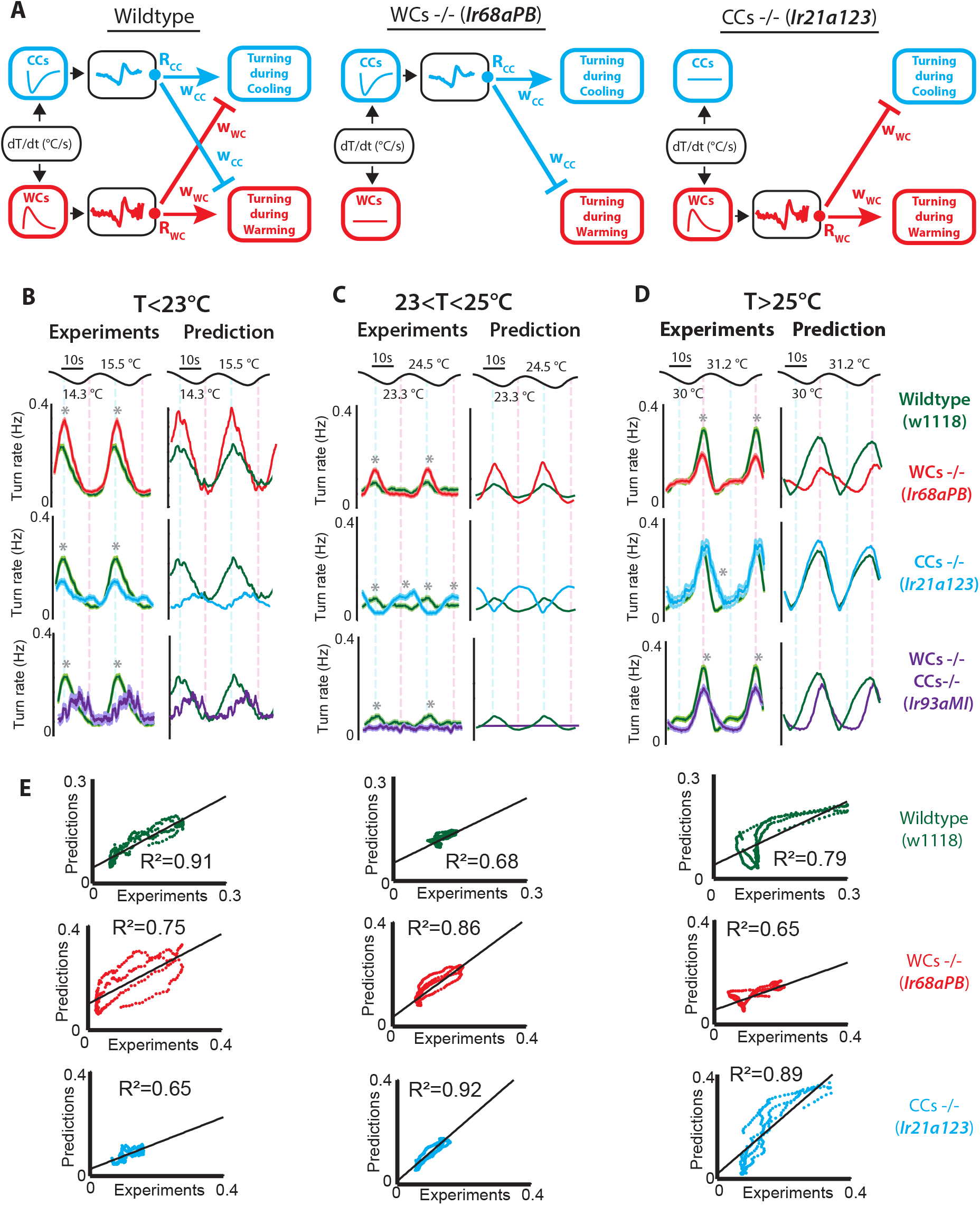
Cross-inhibition computations in mutant larvae. **A**, Schematic representation of the computation underlying thermal homeostasis in wildtype larvae (w^1118^), larvae defective for WCs function (w^1118^; Ir68a^PB^), and larvae defective for CCs function ((w^1118^; Ir21a^123^). **B-D left**, Experimental results of the turning rate response to sinusoidal waves of temperature of wildtype larvae (w^1118^ in green), larvae defective for WCs’ function (w^1118^; Ir68a^PB^ in red), larvae defective for CCs’ function (w^1118^; Ir21a^123^ in cyan), and larvae defective for WCs and CCs function (w^1118^; Ir93a^MI05555^ in purple). Shaded regions are the s.e.m., * indicate different mean turning rate during cooling or warming between the two overlaid turning rate responses in each panel using Chi-squared test with Bonferroni correction (p<0.005). **B-D right**, Model predictions of the turning rate response to a sinusoidal wave of temperature centered at different ambient temperatures. Wildtype larvae’s turning rate in green, larvae defective for WCs’ function in red, larvae defective for CCs’ function in cyan, and larvae defective for WCs and CCs function in purple. **E** Correlation plots between experimental measurements and model predictions for wildtype animals, and mutants defective for WCs or CCs. The experimental results of animals defective for both WCs and CCs are the baseline turning rate of the model. w^1118^ in green, n=80 in **B**, n=50 in **C**, and n=62 in **D**. w^1118^; Ir21a^123^ in cyan, n=45 in **B**, n=54 in **C**, and n=49 in **D**. w^1118^; Ir68a^PB^ in red, n=61 in **B**, n=59 in **C**, and n=55 in **D**. w^1118^; Ir93a^MI05555^ in purple, n=42 in **B**, n=45 in **C**, and n=55 in **D**.

The cross-inhibition computation underlying thermal homeostasis predicts that, below the homeostatic set-point, turning during cooling is strongly stimulated by the CCs and moderately cross-inhibited by the WCs, resulting in a net increase in turning during cooling (Fig6B). Near the set-point, turning during warming and cooling are mutually inhibited by the WCs and CCs, resulting in smaller turning rates during cooling and warming (Fig. 6C). Above the set-point, turning during warming is strongly stimulated by the WCs and moderately cross-inhibited by the CCs, resulting in a net increase in turning during warming (Fig. 6D).

Experiments using mutant larvae confirm all predictions of the cross-inhibition computation. Inactivating WCs (via the *Ir68a^PB^* mutation) reduced turning during warming above the homeostatic set-point (Fig. 6D) and also reduced cross-inhibition of turning during cooling near and below the homeostatic set-point (Fig. 6B, C). Inactivating CCs (via the *Ir21a^123^* mutation) reduced turning during cooling below the homeostatic set-point (Fig. 6B) and reduced cross-inhibition of turning during warming near and above the homeostatic ser-point (Fig. 6C, D). Inactivating both WCs and CCs (by the *Ir93a^MI^* mutation) caused thermal blindness near the set-point (Fig. 6C). However, inactivating both WCs and CCs did not fully abolish behavioral responses far from the homeostatic set-point (Fig. 6B, D). These residual responses are likely due to parallel thermosensory pathways that operate far from the set-point, constituting an additive correction to the computation that integrates WC and CC outputs (Fig. 6B, C, D and Extended Methods).

In addition to predicting the polarity and intensity of behavioral responses, the computation derived here also approximates well the dynamics of the behavioral responses of wildtype and mutant animals, with correlation coefficients between 0.65 and 0.92 (Fig. 6E).

## Neural and behavioral encoding of temperature change speed

The cross-inhibition computation not only explains the polarity and intensity of neural activity and behavior but also the dynamics, enabling us to study how the temporal properties of temperature stimuli are encoded in neural and behavioral dynamics. Intuitively, fast warming above the homeostatic set-point should produces stronger turning responses than slow warming.^29^ Similarly, fast cooling below the homeostatic set-point should produces stronger turning responses than slow cooling. It is not known whether the speed of temperature change is encoded in the neural responses of the sensors or in the transformation from sensory neuron activity to behavior.

We use the cross-inhibition model to formulate hypotheses about how the speeds of warming and cooling are encoded in neural and behavioral responses and test these predictions experimentally. In the cross-inhibition computation, the WCs’ and CCs’ neural responses are passed through linear filters that transform them into the turning rate contributions *R_WC_* and *R_CC_* (Fig. 7A, B top). The linear filters are biphasic in that they have both positive and negative parts. Biphasic filter outputs (*R_WC_* and *R_CC_*) are sensitive to the speed of their inputs (WCs’ and CCs’ neural responses). Thus, *R_WC_* and *R_CC_* are not correlated with WC and CC neural responses; they are correlated with the derivative of WC and CC neural responses (Fig. 7A, B). In consequence, the cross-inhibition computation predicts that even temperature stimuli that produce similar amplitude of WC and CC neural responses can evoke behavioral responses of different magnitude if the speed of the WC and CC neural responses is different. In other words, the speed of temperature change can modulate behavior even if it does not modulate the magnitude of the WC and CC neural responses.

**Figure 7.**
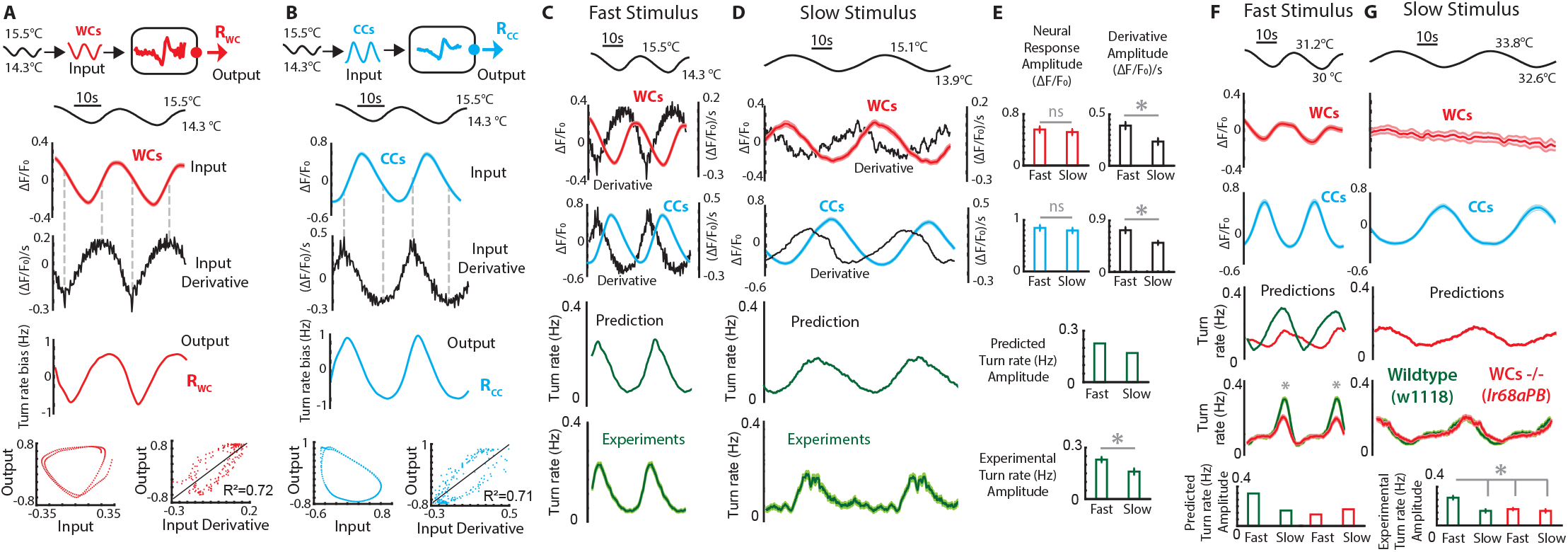
Neural and behavioral encoding of temperature change speed. **A**, **B**, Sinusoidal temperature waves stimulate neural activity changes in the WCs and CCs. The experimentally measured neural activities of WCs and CCs are the input of the cross-inhibition model. In the model, linear filters transform neural activity into the turning rate contributions (*R_WC_* and *R_CC_*). We show in **A** how the WCs activity (the Input) and its derivative (Input Derivative) relate to the turning rate contribution *R_WC_* (Output). The derivative of the WCs activity is strongly correlated with *R_WC_*. We show in **B** how the CCs activity (the Input) and its derivative (Input Derivative) relate to the turning rate contribution *R_CC_* (Output). The derivative of the CCs activity is strongly correlated with *R_CC_*. **C-E**, (First two rows) Neural responses and derivatives of neural responses to slow and fast sinusoidal stimuli centered at low ambient temperatures. The WCs and CCs neural responses to slow and fast sinusoidal fluctuations do not display significantly different amplitudes but the derivatives of the neural responses do display significantly different amplitudes. (Third row) The model predicts different amplitudes of behavioral responses to the slow and fast stimuli. (Fourth row) Following the trend of the model and the derivative of WCs’ and CCs’ activity, experimental results show that turning rate is significantly higher in response to the fast stimulus. **F**, **G**, (First two rows) Neural responses to slow and fast sinusoidal stimuli centered at high ambient temperatures. In this range, the CCs display similar response amplitudes to slow and fast stimuli, but the WCs are unable to detect the slow stimulus. (Third) The model predicts stronger turning rate responses to the fast stimulus, largely due to the contribution of WCs. The green and red curve are identical in response to the slow stimulus. (model predictions without functional WCs in red and wildtype in green). (Fourth and fifth rows) Experimental results follow model predictions, with the contribution of the WCs explaining the difference in turning rates elicited in response to the fast and slow stimuli. In all panels, the * indicate that the peak responses between the brackets are different with Kruskal-Wallis test p<0.01. The grey ‘ns’ indicates no statistically significant difference, and the error bars are the s.e.m.

We tested the model predictions by delivering sinusoidal temperature waves of identical amplitude but different speeds and measuring the WC and CC calcium responses, as well as, the turning rate responses (Fig. 7C, D). Both fast and a slow sinusoidal stimuli evoked WC and CC neural responses with equal amplitude but significantly different response derivatives (Fig. 7C-E top). Because the fast and slow stimuli are centered at a temperature below the set-point, the model predicts an increase in cooling evoked turning in both cases. Even though the WC and CC neural responses have equal amplitudes, the model predicts a stronger behavioral response to the fast stimulus because the turning rate contributions *R_WC_* and *R_CC_* depend on the derivative of the neural responses (Fig. 7C-E). As predicted, wildtype animals also display a stronger cooling avoidance response to the fast stimulus (Fig. 7C-E bottom). The behavioral responses of mutant animals defective for WCs or CCs also follow the model predictions (Supplementary Fig. 9).

Another way in which the speed of warming and cooling can affect behavior is by direct modulation of the WC and CC neural responses magnitude. We tested this at high temperatures, where the WCs are less sensitive to temperature changes. In this temperature range, WCs can detect the fast stimulus but not the slow stimulus (Fig. 7F,G top). Thus, the model predicts that WCs contribute to warming avoidance when exposed to fast stimuli but not when exposed to slow stimuli (Fig. 7F,G). Consistent with the model, the fast stimulus evoked stronger warming avoidance responses, and the mutants defective for WCs function displayed diminished warming avoidance in response to the fast stimulus but identical warming avoidance to wildtype in response to the slow stimulus (Fig. 7F, G).

We conclude that the speed of warming and cooling is encoded in two ways in the cross-inhibition algorithm: (1) in the sensitivity of WCs and CCs to the speed of temperature change, and (2) in the linear filters that transform WC and CC neural responses into the turning contributions *R_WC_* and *R_CC_*. Both types of encoding were experimentally validated using a stimuli that produce WC and CC neural responses of identical amplitude but different speed (Fig. 7C-E), and using stimuli of identical amplitude but different speed that produce different amplitudes of WC neural responses, and consequently, different behavioral responses (Fig. 7F, G).

## Homeostatic temperature control

In many control systems, bidirectional fluctuations from a set-point require bidirectional corrections in opposite directions. We asked whether the homeostatic temperature control system of larval *Drosophila* has this property, and whether near the set-point, the regulation is entirely driven by WCs and CCs.

We used the cross-inhibition model to identify the experimental conditions required for bidirectional control. In the cross-inhibition model, *w_WC_* obtains larger values at higher ambient temperature, and *w_CC_* obtains higher values at lower temperatures. In order, to use the model to predict behavioral responses at all ambient temperatures, we fitted linear functions that relate ambient temperatures with the values of *w_WC_* and *w_CC_* (Fig. 8A, B). Then, we used fast and slow sinusoidal stimuli as inputs to the model and calculated the predicted turning rates for all ambient temperatures (Fig. 8C).

**Figure 8.**
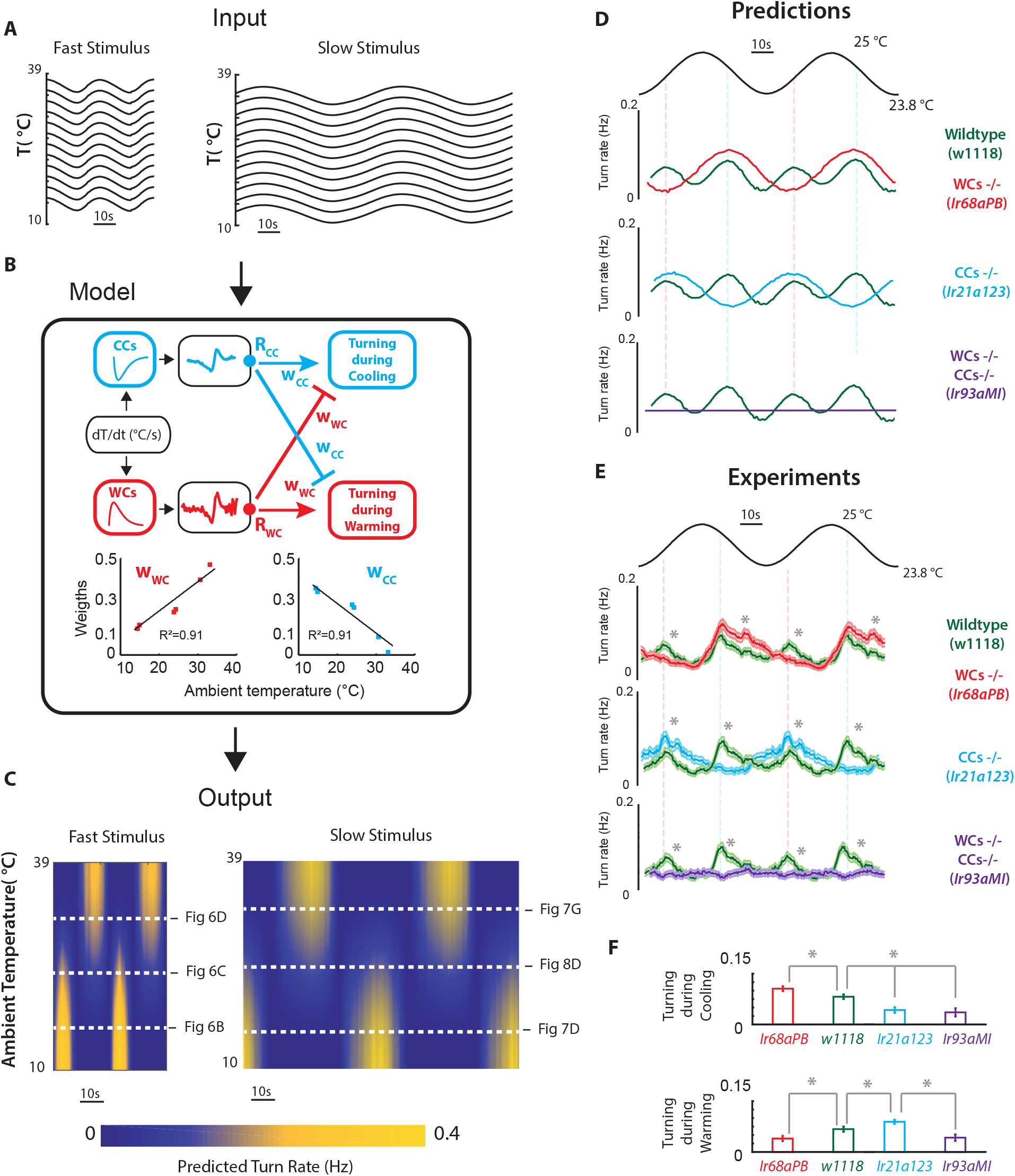
Homeostatic temperature control. **A**, Sinusoidal temperature stimuli at all ambient temperatures can be used as an input to the model. **B**, In order to predict behavioral responses at all ambient temperatures, we use linear filters to predict neural responses and we fit linear functions that relate the weights (*w_WC_* and *w_CC_*) to ambient temperature. **C**, Using the Cross-Inhibition Model we predict the behavioral responses driven by the WCs and CCs pathway for stimuli at all ambient temperatures. **D**, A stimulus centered at the homeostatic set-point is predicted to drive turning increases during both cooling and warming. The turning during warming is driven by the WCs and cross-inhibited by the CCs. The turning during cooling is driven by the CCs and cross-inhibited by the WCs. **E**, **F**, Experimental results validate all model predictions. In all panels, the * indicate that the peak responses between the brackets are different with Kruskal-Wallis test p<0.01. All error bars and shaded regions represent the standard error of the mean (s.e.m.).

The model predictions reveal a temperature range around 24.5°C where bidirectional control occurs. Bidirectional control was not observed in Fig. 6C, for example, because the sinusoidal temperature wave did not go above 24.5°C. Around 24.5°C, wildtype animals are predicted to increase their turning rate in response to both cooling and warming (Fig. 8D green line). Mutants defective for WCs function are predicted to display exacerbated turning during cooling and baseline turning during warming (Fig. 8D red line). Mutants defective for CCs function are predicted to display exacerbated turning during warming and baseline turning during cooling (Fig. 8D cyan line). Mutants defective for both CCs and WCs are predicted to display no behavioral responses (Fig. 8D purple line).

Experimental results validate all model predictions near the set-point: Wildype animals have increased turning during warming and cooling; mutants defective for WCs function have exacerbated turning during cooling and do not modulate turning during warming; mutants defective for CCs function have exacerbated turning during warming and do not modulate turning during cooling; without WCs and CCs there are no behavioral responses (Fig. 8E, F).

We conclude that both WCs and CCs are required for homeostatic temperature control. Missing one type of sensor results in avoidance to only warming or cooling and an inability to stay at the homeostatic set-point. Cross-inhibition scales the intensity of the avoidance response to the adversity of the stimulus.

## Discussion

In thermosensation, often one type of sensor (cooling or warming) is studied without its counterpart, making our understanding of the computations underlying thermal homeostasis incomplete.^1, 18, 30^ The justification for studying thermosensory cells function without their counterparts has been largely supported by the labeled lines hypothesis, in which cells activated by cooling exclusively regulate mechanisms that counteract cooling and cells activated by warming exclusively regulate mechanisms that counteract warming.^31–33^ In this view, the outputs of warming and cooling cells do not necessarily integrate to shape thermoregulatory responses. However, recent work in adult *Drosophila* has shown that specific interneurons explicitly integrate cooling and warming stimuli,^34, 35^ suggesting that cooling and warming pathways may combine to shape behavior. In mammals, cooling cells may cross-modulate mechanisms that counteract warming and warming cells may cross-modulate mechanisms that counteract cooling, but these mechanisms remain unexplored.^2, 36^

Here, we discovered the larval *Drosophila* warming cells (WCs) and studying them together with their counterparts, the cooling cells (CCs), we uncovered the computation underlying thermal homeostasis. In this computation, WCs and CCs outputs combine via cross-inhibition to drive behavioral responses to both cooling and warming stimuli.

### Encoding of thermosensory polarity by ancestral ionotropic receptors

The new set of WCs we presented here share anatomical and genetic similarities with the CCs. Each sensor requires a distinct but partly overlapping set of Ionotropic Receptors to detect temperature changes. The WCs require Ir68a, Ir93a, and Ir25a to detect warming. The CCs require Ir21a, Ir93a, and Ir25a to detect cooling. The opposed thermosensitive polarity of the WCs and CCs is encoded in their heteromeric expression of sets of Ir receptors. These Ionotropic Receptors are conserved across insects^37^ and have homologs in the disease vector mosquitoes *Anopheles gambiae* and *Aedes aegypti*, which use temperature cues to identify human hosts.^38, 39^ The combinatorial use of ionotropic receptors may be a widely used mechanism to shape the sign and sensitivity of thermosensory responses in different insects.

### Relevance of synchronous and opponent sensors encoding

Many sensory modalities use opponent sensors to encode environmental stimuli including photosensation, hygrosensation and thermosensation.^1, 18, 30^ The larval WCs and CCs respond to temperature changes with opposite polarity, but with symmetric temporal dynamics. Calcium imaging revealed that WCs and CCs share the same peak times and adaptation times to a variety of sensory stimuli. When we exposed freely moving animals to optogenetic white noise stimulation of WCs or CCs and conducted reverse-correlation analysis, we also uncovered identical transformations that convert WC and CC activity into synchronous changes in behavior. This synchrony facilitates the integration of the output of WCs and CCs by downstream circuits. In particular, synchrony makes it possible to use linear combinations of WC and CC outputs to determine behavioral responses. The use of sensors with opposed polarity has been proposed to increase the optimality of information encoding.^40^ Our study underscores a potential advantage in signal processing of having synchronous sensors with opposed polarity.

### Cross-inhibition computations in biology

We found that larval *Drosophila* uses a flexible cross-inhibitory computation to achieve thermal homeostasis. Above 24°C warming is unfavorable because it carries the animal further from the homeostatic set-point. In this temperature range, avoidance responses during warming are strongly stimulated by WCs and moderately cross-inhibited by CCs. Below 24°C, cooling is unfavorable. In this temperature range, avoidance responses during cooling are strongly stimulated by CCs and moderately cross-inhibited by WCs. Near the homeostatic set-point, balanced cross-inhibition from WCs and CCs suppresses avoidance responses.

Cross-inhibition is prevalent in perceptual-choice models. In these models, cross-inhibition between competing groups of neurons often enhances accuracy in decision-making. For example, in the Usher-McClelland model of primate decision-making, different neuron groups are used to represent different choices.^41^ These neurons mutually cross-inhibit their output pathways. The most strongly activated group that represents a specific choice thus biases the decisions towards one outcome by suppressing all others.^41, 42^ In larval *Drosophila* thermoregulation, the choice is whether to avoid cooling or warming: at high temperatures, warming should be avoided; at low temperatures cooling should be avoided. At all temperatures, however, the CCs are always more active during cooling and the WCs are always more active during warming. Thus, any cross-inhibition in the outputs of the WCs and CCs has to be flexible for these neurons to contribute differently to behavior in different contexts. Unlike in the Usher-McClelland model, in the case of larval *Drosophila* thermoregulation, flexibility is encoded in the ambient temperature-dependent weights of the WC and CC contributions to behavior, and not in the WC and CC neural activity.

### Consequences for thermal homeostasis

In mammals, prevailing models of thermoregulation propose that signals from cooling and warming cells are integrated in the preoptic area of the hypothalamus.^2^ GABAergic and Glutamatergic neurons are proposed to play a role in modulation of hypothalamic warming cells.^2–4^ However, the computations driving thermal homeostasis in mammals remain obscure and are challenging to dissect because overlapping autonomic and behavioral mechanisms contribute to thermoregulation, making the output of the computation multidimensional.

Because poikilotherms strictly use behavior for thermoregulation, measurements of behavior constitute the full output of the thermoregulatory computation. Like mammals, *Drosophila* integrates the outputs of bidirectional sensors of cooling and warming to regulate body temperature. The computation underlying thermal homeostasis in the larva may represent a general means of maintaining a set-point using opponent sensors.

### Significance for Systems Neuroscience

Many poikilotherms, including cave beetles^6^ and python vipers,^43^ have neural and behavioral responses to temperature changes that are as robust as the ones displayed by larval *Drosophila*. The significance of understanding the computations underlying thermal homeostasis in *Drosophila melanogaster* is that, unlike other poikilotherms, *Drosophila*’s genetic accessibility allows to manipulate with precision the individual receptors and neurons underlying thermal homeostasis. In addition, the recent advances in larval *Drosophila* connectomics^44^ indicate that the neural implementation of the computation we discovered here, can potentially be determined by tracing the synaptic connections of the neurons downstream from the WCs and CCs, and measuring neural activity of those neurons.

### Consequences for other homeostatic control systems

Homeostatic control is pervasive in biology, including homeostatic control of synaptic plasticity in connections between neurons,^45^ homeostatic control of cardiac output,^46^ and homeostatic control of glucose.^47^ All homeostatic processes regulate a physiological variable near an optimal set-point. We have identified the control system for homeostatic thermal regulation in larval *Drosophila*. We have fully determined the computation that integrates the outputs of warming and cooling cells, and established how this computation leads to control over the homeostatic variable. Our analysis establishes a framework for gaining computational insight into a homeostatic control system.

## Methods

### Fly husbandry

Flies were raised at constant temperature (22 °C) and 50% humidity on standard cornmeal agar-based medium. For experiments with larvae, adult *Drosophila melanogaster* were transferred to collection cages (Genesee Scientific). One end of the cage held a grape juice agar plate and fresh yeast paste. Flies laid eggs on the agar plate for 2 days when the plate was removed to collect larvae. For all experiments, early second instar larvae were selected based on spiracle development using a dissection microscope.

### Genotypes

The genotypes of fly stocks used in this study:

**Effectors:** UAS-GCaMP6m in the second chromosome: w[1118]; P[y[+t7.7] w[+mC]=20XUAS-IVS-GCaMP6m]attP40 (from BDSC 42748). UAS-GCaMP6m in the third chromosome: w[1118]; PBacy[+mDint2] w[+mC]=20XUAS-IVS-GCaMP6mVK00005 (from BDSC 42750). UAS-GFP: w[*]; P[y[+t7.7] w[+mC]=10XUAS-IVS-mCD8::GFP]attP40 (from BDSC 32186). UAS-Ir68a and UAS-Ir93a from.^23^ UAS-Ir25a from.^48^ UAS-CsChrimson: w[1118]; Py[+t7.7] w[+mC]=20XUAS-IVS-CsChrimson.mVenusattP2 (from BDSC 55136).
**Gal4-Drivers:** pebbled-Gal4 from.^11^ w[1118];P[Ir68a-Gal4]attp2 backcrossed from,^23^ and w[1118]; Py[+t7.7] w[+mC]=GMR11F02-GAL4attP2 from BDSC 49828.
**Mutants:** Ir68a^PB^ from,^23^ Ir93a^MI05555^ from,^18^ Ir21a^123^ from,^17^ Ir25a^2^ from,^25^ Ir25a-BAC from ^2^.

### Confocal microscopy

All fluorescence imaging was performed using a spinning-disk confocal setup using a 60x 1.2-N.A water immersion objective (Nikon Instruments LV100; Andor). During functional imaging in response to temperature changes, thermal expansion of the objective lens was compensated using a piezoelectric element.^1^ For each experiment, larvae were washed with water and partially immobilized under a cover slip.^1^ The microscope stage was temperature controlled using a Peltier thermoelectric actuator (Custom Thermoelectric) controlled with an H-bridged power driver and a two-degrees-of-freedom PID control algorithm. This algorithm was implemented using a PID control module (Accuthermo Technologies) operated by custom code written in LabView (See Extended Methods). The Peltier element was cooled by flowing antifreeze through an attached water block. The antifreeze was kept at 8-10°C using a VWR chiller.

### Temperature-controlled behavioral apparatus

The temperatre controlled behavioral apparatus was operated inside a dark enclosure to prevent any light from causing phototactic artifacts. The behavioral arena was mounted on vibration-damping legs to eliminate mechanical artifacts. Dark-field illumination was provided with custom-built infrared LED bars (*λ* 850nm) operated with 10% pulse width modulation to avoid heating artifacts. The behavioral arena was temperature controlled with four Peltier thermoelectric actuators (Custom Thermoelectric) controlled with an H-bridged power driver and a two-degrees-of-freedom PID control algorithm. This algorithm was implemented with PID control modules (Accuthermo Technologies) and custom code written in LabView (See Extended Methods). Feedback signals for PID control were from thermo-couples located in the behavioral arena as well as on the agar.

During behavioral experiments 15-18 larvae crawled freely for 20 min on 10×10 cm agar squares with 4mm thickness. These surfaces contained 2% agar and 0.1% activated charcoal (Sigma-aldrich). Charcoal increases visual contrast when imaging. We captured movies using a CCD camera (Mightex) with a long-pass infrared filter (740nm) at 4 fps.

### Optogenetic behavioral apparatus

For optogenetic experiments, animals were reared in cages with grape juice plates with a mixture of 0.18 grams of yeast and 400 uL of 0.5mM all-trans-retinal (ATR). The cages were kept in complete darkness until the experiment. The setup for optogenetic behavioral experiments is described elsewhere (^27^). In brief, optogenetic light stimulation was provided by a custom built LED matrix (SMD 5050 flexible LED strip lights, 12V DC, *λ* 625nm). Optogenetic stimulation was controlled with an H bridge driver and custom code written for a LabJack U3 controller. Light intensity was controlled via pulse-width-modulation at 500kHz. Optogenetic stimulation was synchronized with image acquisition. Dark-field illumination was provided using custom-built infrared LED light bars (*λ* = 850nm). The wavelength of infrared illumination was chosen to avoid interference with the red LED illumination for optogenetic stimulation. Infrared LEDs for dark field illumination were mounted using opto-mechanical elements to adjust the angle with respect to the behavioral arena, avoiding larval ‘shadows’ that lowered the efficiency of data acquisition. The red LEDs were connected in parallel to produce uniform illumination. We verified uniform light intensity at 1.5 W/m^2^ 0.02. The behavioral arena was 22×22 cm and used the same agar composition as for temperature-controlled behavior experiments. In each experiment, 25-30 larvae were used and their movements were recorded with a CCD camera with a long-pass infrared filter (740nm) at 4 fps. Temperature was controlled in the same way as for thermoregulation behavior experiments.

### Focused ion beam scanning electron microscopy (FIB-SEM)

For FIB-SEM serial sectioning, second third instar wild type Canton S larvae were used. After rinsing in PBS, the anterior half of the larva was incubated in fixative (2% formaldehyde with 2.5% glutardialdehyde in 0.1 M Na-cacodylate buffer, pH 7.4, Sigma-Aldrich, Germany,) for 30–90 min. Then, the head region was cut off and incubated in fresh fixative for 90 min. Samples were washed in Na-cacodylate, followed by post-fixation in 1% osmium tetroxide (SERVA Electrophoresis GmbH, Germany) for 2 hr at 48°C in the dark. En bloc staining was carried out with 1% uranyl acetate and 1% phosphotungstic acid in 70% EtOH in the dark over night before continuing the alcohol dehydration the next day. Samples were transferred to propylene oxide before embedded in Spurr (Plano,GmbH, Germany) using ascending Spurr concentrations diluted in propylene oxide for optimal tissue infiltration. Polymerization was carried out at 65°C for 72 hr. Blocks were trimmed using an Ultracut UCT microtome (Leica, Germany), mounted on conventional SEM stubs, and sputtered with 80–100 nm platinum. FIB-SEM serial sectioning was carried out using a FESEM Auriga Crossbeam workstation (Zeiss, Germany). FIB fine milling was carried out with 500 pA.

### Behavioral quantification

Behavior was pre-processed using MAGAT Analyzer (https://github.com/samuellab/MAGATAnalyzer). Every larva image was used to calculate its mid-line. Each mid-line was then segmented in 11 points. Eight behavioral parameters where calculated from the body contour and segmented mid-line: speed, crab-speed, spine length, direction of motion, forward/backward crawling bias, head turn, head angular speed, and area of the larvae body (see extended methods for details). The time traces of these behavioral parameters over one period of a temperature sine wave stimulus were used to build an interpoint dissimilarity matrix, followed by multidimensional scaling, dimension selection, and an iterative denoising trees algorithm to classify larvae motor sequences in response to temperature fluctuations. This procedure was implemented following.^49^ See the details of the calculations in the Extended Methods.

### Computations underlying thermal homeostasis

One component of the thermoregulatory computations is the filter that transforms the neural activity of CCs or WCs into behavioral responses. We estimated these filters by combining results from our calcium imaging experiments and optogenetic behavior experiments (see Extended Methods and^27^). In brief, the normalized measured activity responses of WCs and CCs (measured by calcium imaging) were convolved with the linear filters that convert WC and CC activities into behavioral responses (measured using optogenetics and quantitative behavioral analysis). The result of these convolutions were weighted to reflect the contribution of each sensor type to behavioral response as follows:

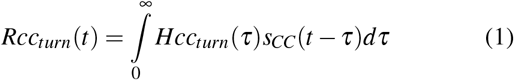

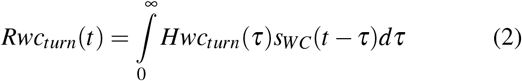

Where *Hwc_turn_* and *Hcc_turn_* are the convolution kernels for the WCs and CCs, respectively. Each kernel is computed from the signal history of the WCs (*s_WC_*(*t* − *τ*)) and CCs (*s_CC_*(*t* − *τ*)). The turning rates calculated from equations (1) and (2) are linearly combined with scalar weights, *w_CC_* and *w_WC_*, for all models to obtain the predicted turning rate *R_turn_*(*t*) as follows:

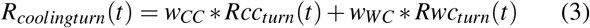

In the *Labeled Lines Model (LLM)*, turning during cooling can only be modulated by the CCs and turning during warming can only be modulated by the WCs. These conditions amount to a rectification of the *Rwc_turn_* during cooling and *Rcc_turn_* during warming.

In the *Cross-Inhibition Model (C-IM)*, turning is controlled by both WCs and CCs. The contributions of WCs and CCs (*Rwc_turn_* and *Rcc_turn_*) to behavior are allowed to take negative values and therefore do not require a transformation to positive values nor a saturation before the linear combination. Turning rates cannot be negative though, so the result of the linear combination is transformed with a linear function into non-negative values.

In the *Cross-Activation Model (C-AM)* turning is controlled by both WCs and CCs. The contributions of WCs and CCs to behavior (*Rwc_turn_* and *Rcc_turn_*) are transformed with a linear function to have non-negative values. The result of the linear combination will be always positive so it does not need to be transformed again.

In all cases, the values of the weights (*w_CC_* and *w_WC_*) are linearly regressed to match the amplitude of the wildtype animal responses. See the Extended Methods for a detailed derivation of the models.

## Supporting information

Supplementary Material Part 1

Supplementary Material Part 2

## ACKNOWLEDGEMENTS

We acknowledge members of the Samuel and Garrity labs for helpful discussions and comments on the manuscript. We thank Rachel Wilson, Sandeep Robert Datta, and Alex Schier for helpful discussions and advice during the development of the project. We acknowledge the Bloomington *Drosophila* Stock Center (NIH P40OD018537). L.H-N., G.B., P.G. and A.D.T.S. were supported by the NIH R01 GM130842-01 grant. A.S.T., V.R. and A.R., were supported by the Deutsche Forschungsgemeinschaft (TH1584/3-1, TH1584/6-1 and TH1584/7-1).

## AUTHOR CONTRIBUTIONS

L.H-N. developed the experimental techniques for temperature control and optogenetics, wrote the software for behavioral classification, analyzed data, performed behavioral experiments, functional and anatomical imaging, and optogenetic experiments, and derived the thermoregulation computational models. A.C. performed behavioral experiments, functional imaging, and optogenetic experiments. G.B. conducted immunostaining and anatomy experiments. V.R., A.R. and A.S.T. contributed the Electron Microscopy dataset and did the EM reconstruction. M.K. provided reagents, preliminary data, and technical advice. L. H-N., P.G. and A.D.T.S. designed the study, interpreted the results, and wrote the manuscript with feedback from all authors.

