## Extended Methods for "Synchronous and opponent thermosensors use flexible cross-inhibition to orchestrate thermal homeostasis"

#### CNMF sensory neuron's analysis

We used a constrained non-negative matrix factorization (CNMF) framework[12] to measure the responses of all larval head sensory neurons to temperature fluctuations. We used pebbled-Gal4 to label all larval head sensory neurons with GCaMP6m[16,19] (Fig. 1A) . These neurons are housed in three ganglions: the dorsal organ ganglion (DOG), the terminal organ ganglion (TOG) (Fig. 1A), and the smaller ventral organ ganglion (not shown).[16] The fluorescent baseline of different sensory neurons differs in as much as two orders of magnitude, so visual inspection is insufficient in identifying simultaneous neural responses. After denoising, normalizing, and identifying the regions of interest (Fig. 1B), we were able to identify an average of 43 regions of interest (ROIs) per animal: 38 of those ROIs did not present activity correlated with the stimulus (Supplementary Fig. 1C); three cells responded to cooling (Supplementary Fig. 1A), which are the cooling cells (CCs) previously discovered in;[1] and, notably, two cells responded during warming (Supplementary Fig. 1B). We refer to these two cells as the warming cells (WCs).

#### Ionotropic Receptors expression in the Dorsal Organ Ganglion

A previous study used RNA-sequencing to find the Ionotropic Receptors expressed in the larva's Dorsal Organ Ganglion (DOG).[53] This study found that Ir25a, Ir47a, Ir92a, Ir93a, Ir21a, Ir68a, Ir62a, and Ir76b are consistently expressed in the DOG. A second study, generated Ir-Gal4 lines and verified expression in the DOG, narrowing down the search further to Ir25a, Ir93a, Ir76b, Ir68a, Ir21a, and Ir92a.[48] From these 6 receptors, Ir25a, Ir93a and Ir21a are expressed in the cooling cells (CCs) and required for cooling sensing. We tested the remaining 3 receptors, Ir76b, Ir92a, and Ir68a. We did not find any fluorescence in Ir76b at the second instar stage of larva development. Ir92a is expressed in one neuron in the DOG. We expressed GCaMP6m in Ir92a-Gal4 and exposed those larvae to temperature fluctuations but this neuron did not display thermosensitivity. Finally, Ir68a-Gal4 labeled the two warming cells (WCs) and did not show expression in other parts of the larva (Supplementary Fig. 2A-C).

#### Gal4 lines expression patterns

An important control for optogenetic experiments is the specificity of the Gal4 lines driving expression of CsChrimson. In a previous study,[1] R11F02-Gal4 showed expression in the CCs. We tested expression patterns using Ir68a-Gal4, Ir21a-Gal4 and R11F02-Gal4 to drive CD8-GFP expression in orco-RFP background larvae. We found that Ir68a-Gal4, R11F02-Gal4 but not Ir21a-Gal4 drivers were very specific for WCs and CCs respectively at the early second instar stage. Ir68a is specific in the entire animal (Supplementary Fig. 3a), while 11F02 Gal4 is also highly specific with occasional expression in an additional pharyngeal neuron (Supplementary Fig. 3b). The olfactory dome, the intestines, and the tail are auto-fluorescent because they are fluorescent in either the green or red channels (Supplementary Fig. 3 insets).

#### Scanning Electron Microscopy

For SEM, second instar Canton S wild type larvae were prepared. After rinsing the larvae in PBS, they were bathed in hot water (90°C) for 1.5–2 min. Subsequent fixation was carried out in 2.5% glutardialdehyde (Agar Scientific, UK) buffered in Na-cacodylate buffer at 48°C. After 1/2 hr, the anterior first third of larvae was cut off and put back in fresh fixative for additional 3 hr at 48°C. After fixation, samples were washed in Na-cacodylate, followed by post-fixation in 1% osmium tetroxide (SERVA Electrophoresis GmbH, Germany) for 2 hr at 48°C in the dark. After additional washing steps in Na-cacodylate, samples were dehydrated in ascending ethanol concentrations at 48°C. Then, samples were dried via CO2 (Bal-Tec CPD 030, Liechtenstein). Specimens were coated with gold-paladium or platinum. Samples were examined in a FESEM Auriga Crossbeam workstation (Zeiss, Germany).

In the EM volume, the ORNs were easy to identify because of their stereotyped dendritic bundles. Thus, to identify the WCs and CCs we considered the relative location of these cells to the olfactory receptor neurons (ORNs), and the shapes of the outer dendritic processes observed via confocal microscopy in both WCs and CCs. The A-type CCs are more dorsal than the B-type CC and are the most medial and lateral CCs.

#### Controlling Temperature Waveforms

To study thermotaxis we used a temperature-controlled platform (Fig. 3A).[1] We recorded larvae movement with a CCD camera and provided dark-field illumination with low intensity infrared LEDs. The stage temperature can be heated with different waveforms using four thermoelectric elements that use the Peltier effect to either heat or cool the stage. Water blocks are used as heat exchangers: coolant liquid flows propelled by a system of pumps in series cooling down the thermoelectric elements. The coolant liquid is kept at 5°C in a VWR chiller. Our control strategy consisted in measuring the input/output of the system (the input is the desired temperature wave and the output the resulting temperature wave) and using those measurements to fit an autoregressive moving average with exogenous input (ARMAX) model to the temperature control system[55] (Supplementary Fig. 7). This was implemented using the Systems Identification Toolbox in MATLAB. The ARMAX model is then used to run simulations of a second degree of freedom of a PID (Proportional, Integral, Derivative) control[56] of the system and we select the control parameter combinations that produce the closest signal to the one used for microscopy experiments.

The 1-DOF PID controller has the form of the following equation:

$$C(s) = K_p(1 + \frac{1}{T_I s} + T_D s) \quad (4)$$

Where $C(s)$ is the transfer function of the 1-DOF PID controller. The Laplace transform was used to transform the function from the time real variable $t$(time) to the complex variable $s$ (frequency). The constants $K_p$(Proportional component), $T_I$(Integral component), and $T_D$(Derivative controller), are tuned to eliminate overshooting of the system and guarantee convergence between the desired set-point and the experimental set-point. The second degree of freedom of the PID controller (also called filter compensator) uses additional parameters with proportional, integral, and derivative action to improve control. The filter compensator transfer function ($F(s)$) depends on the original PID parameters and on the new parameters $\gamma$, $\alpha$, and $\beta$:

$$F(s) = \frac{\gamma + (1-\alpha)T_I s + (1-\beta)T_I T_D s^2}{1 + T_I s + T_I T_D s^2} \quad (5)$$

The new parameters $\gamma$, $\alpha$, and $\beta$ are tested recursively with the ARMAX model of the temperature control apparatus. We vary the three parameters from 0 to 1 and run simulations with thousands of different parameter combinations until the result fits the desired signal with $R^2 > 0.9$. Then we use these parameters in custom code written in LabVIEW as the input of the temperature control system (https://github.com/LuisM6/ThermotaxisCrossInhibition2020). Using this temperature control technique enabled us to produce identical signals for behavioral measurements and calcium imaging (Supplementary Fig. 7).

### Unsupervised learning of behavior sequences

Clustering techniques are easy to use to classify elements based on features with scalar values. These methods have been used to describe behaviors based on the mean of animal posture and trajectory parameters over the time of the experiment.[57] These approaches fail to describe appropriately the sequences of motor outputs used by the animal during the experiment because they average over time. A more complex computational task is to classify behaviors based on features represented by time series. We implemented a multivariate time-series unsupervised classification algorithm similar to the one presented in[49] to distinguish behaviors based on sequences of motor outputs described by the time series of animal posture and trajectory features.

Our behavioral processing pipelines has three main parts. First we collected movies of 20 minutes of larvae exposed to identical sinusoidal waves of temperature centered at 15.2°C (cyan curve), 23.8°C (black curve), and 30.6°C (red curve) (Supplementary Fig. 8A). These movies were processed using MAGAT analyzer (https://github.com/samuellab/MAGATAnalyzer) to obtain the coordinates of each larva's body contour (in red). From the contour we calculate the center-line of the larvae and segment it in 11 points (in yellow). Assuming, larvae spend more time moving forward than backwards the head and tail were labeled (head in green and tail in blue) (Supplementary Fig. 8A). Using this coordinate system for the larva's body, we calculate the time-series of features that describe behavior (https://github.com/LuisM6/ThermotaxisCrossInhibition2020) :

• Speed: Magnitude of the velocity vector of the centroid of the larva. This quantity is normalized by the mean speed in the absence of stimulus.
• Crabspeed: Magnitude of the projection of the velocity vector projection in the axis that is aligned with the best fit of a unit vector to the larva spine.
• Spine length: Euclidean distance normalized to the mean length in the absence of stimulus.
• Direction of motion: One minus the product of the unit vectors of the current and previous path segments. It is zero for straight motion and -2 for perfect reversal.
• Forward/Backward crawling bias: When the projection of the velocity vector in the tail to head vector is positive (the animal moves in the headward direction) the crawling bias is 1; when the projection is negative -1, and 0 when the animal is not moving.
• Head turn: Angle in radians between the head and the spine.
• Head sweep angular speed (rad/s): Angular speed of the head with respect to the spine.
• Area of the larva body: Area of the larva body normalized by the mean area measured in the absence of stimulus.

This parameter selection has proven effective to describe larvae behavior.[49] We normalize speed and crabspeed to avoid dependence on larval size.

The time-series of these features are the input to the unsupervised behavioral classifier. The algorithm consists in first building an inter-point dissimilarity matrix, which is the input for Multidimensional scaling. Later we conduct dimensionality selection and finally we run the iterative denoising trees clustering algorithm recursively until we reach convergence. The construction of the inter-point dissimilarity matrix, the multidimensional scaling and selection are in https://github.com/LuisM6/ThermotaxisCrossInhibition2020, the output of these scripts is then imported in R and is the input for the interative denoising trees clustering algorithm from.[49] The output of the clustering algorithm is later processed to identify the time traces of the behavioral features of each cluster in MATLAB.

The larva is believed to use turning rate to bias its random walk during navigation.[1,26,27] Analyzing behavior in response to other sensory modalities with this technique, we found, for example, that in addition to turning, certain stimuli can drive reversals and speed modulation (unpublished). Thus, we analyzed the thermotaxis data with this same algorithm to explore this possibility. We did not find evidence of reversals or speed modulation in thermotaxis (Supplementary Fig. 8B). Instead, we found that only turning during cooling and turning during warming are are sampled from different distributions at different temperature baselines in wildtype and mutant larvae (Steel-Dwass test p<0.01). This finding is consistent with previous studies[1,26,27] and is the first unsupervised (not concluded by human observation) proof of turning rate being the main driver of temperature-evoked behavior in larvae. Thus, we use turning rate as the behavioral readout of behavior.

While iterative denoising trees helps us classify the motor sequences used by larvae to respond to sensory stimuli, it could only classify turns in four time bins: early cooling (behavioral sequence 9), late cooling (behavioral sequence 10), early warming (behavioral sequence 11), and late warming (behavioral sequence 12) (Supplementary Fig. 8C). Because we want to understand how temporal patterns of neural activity are transformed into behavior, we recalculate turning rate timing with more precision, using the machine vision sequence proposed in.[58] This algorithm flags a turn when the angle between the larvae neck (first 3 of 11 segments) forms an angle of more than 20° with the spine (last 8 segments of the 11) and when the larvae has reached a local speed minimum.

### Perceptual choice model of thermotaxis

Understanding how thermal stimuli is transformed into behavior requires multiple measurements and the quantification of signal transduction and sensorimotor transformations. Here we detail the different steps of our model.

### Sensory transduction:

In the first layer of the model we want to quantify how temperature changes are encoded in the WCs and CCs. We used Gal4 drivers to express GCaMP6m in the WCs and CCs, and exposed larvae to sine waves of temperature with different baselines (15, 24, and 32°C) while doing volumetric imaging with a spinning disk confocal microscope. Each neuron's fluorescence response ($\Delta F = F - F_o$) was normalized by its baseline fluorescence ($S = \Delta F/F_o = (F - F_o)/F_o$) at the temperature baseline of the experiment in the absence of temperature fluctuations. Where $S$ is the response reported in Fig. 2 and all other figures involving calcium imaging.

Fluorescence measurements and baseline fluorescence, both depend on the level of expression of GCaMP6m. For example, having two copies of UAS-GCaMP6m or two copies of the Gal4 driver as opposed to one can generate differences in the amplitude of our measurements. To remove this artifact from the model we calculate the normalized responses $S_{norm}$ by further normalizing the calcium responses $S$ by $S_{max}$ ($S_{norm} = S/S_{max}$), where $S_{max}$ is the maximum absolute value of the measured calcium responses $S$ to fast and slow sine waves of each sensor type. This guarantees that all the normalized neural responses are between 1 and -1, and captures the timing of the response and the relative amplitudes for different baselines and speeds of the sinusoidal stimulus.

Because the timing of the responses of A and B-type sensors occur in the same timescales (Supplementary Fig. 10), and because all the CCs form one glomerulus in the antennal lobe,[1] we consider them both as a unit in the model. The small differences in the A and B-type CCs' response amplitude do not affect the conclusions of the model. We used the A-type CCs responses for the calculations we show in Fig. 3; however, using the B-type CCs produces equivalent results.

### Reverse correlation using optogenetics:

In[27] we introduced a method to quantify larvae navigation dynamics using optogenetics and linear filters. Here we use the same technique. We expressed CsChrimson in either CCs and WCs, and exposed freely moving larvae to a Gaussian White noise of red light intensity with mean 1.5 W/$m^2$ and variance 1. White noise frequency is synchronized with the camera frame rate as described in the Methods section. The entire arena is subject to spatially-uniform fluctuations of light intensity with +/- 0.02 W/$m^2$. White noise allows us to recover linear filters because the average optogenetic stimulus history preceding a turn is proportional to the linear filter of a turning event.

We model the larva as a linear transducer. By definition, the probability per time unit of initiating turn, in other words, the turning rate $R_{turn}(t)$, is a weighted sum of stimulus history, $s(t)$:

$$R_{turn}(t) = \int_0^\infty H_{turn}(\tau)s(t-\tau)d\tau \tag{6}$$

Where the integral is the filter and $H_{turn}$ is the kernel.

When the stimulus is a Dirac delta function ($\delta(t)$), the response is a direct measurement of the kernel:

$$R_{turn}(t) = \int_0^\infty H_{turn}(\tau)\delta(t-\tau)d\tau = H_{turn}(\tau) \tag{7}$$

Alternatively, the first order filter can be recovered by measuring the response to a white process as Gaussian white noise. The linear filter relates the cross correlation of input and output $C_{rs}$) and the autocorrelation of the input ($C_{ss}$) by:

$$C_{rs}(t) = \int_0^\infty h(\tau)C_{ss}(t-\tau)d\tau \tag{8}$$

For a Gaussian white noise process, as the one used here, the autocorrelation function ($C_{ss}$) depends on the variance and the Dirac delta function ($\sigma^2\delta(t)$), our variance was chosen to be 1, therefore:

$$C_{rs}(t) = \int_0^\infty H(\tau)\delta(t-\tau)d\tau = H(t) \tag{9}$$

Thus, the cross-correlation of input and output represents a measurement of the linear filter. In our analysis the relevant events in the output is the turning rate. The average optogenetic activation signal that precedes each turning events is called the event-triggered average ($M$) and can be written as shown below:

$$M(\tau) = \frac{1}{n}\int_0^T R_{turn}(t)s(t-\tau)dt \tag{10}$$

Where $n$ is the average number of turns per trial, and $T$ is the duration of each trial.

The cross-correlation function ($C_{rs}$) can be written:

$$C_{rs}(\tau) = \frac{1}{T}\int_0^T R_{turn}(t)s(t+\tau)dt \tag{11}$$

From equations (11), (12) and (13):

$$H(\tau) = C_{rs}(\tau) = \frac{n}{T}M(-\tau) \tag{12}$$

Thus, we estimated our linear filters by measuring the event-triggered average, and multiplying by the average number of events in one trial divided by the duration of the trial.

The most valuable information of a kernel is encoded in its timing because the amplitude of it can depend on the expression level of Gal4 drivers, thus kernels are normalized in the computational model. A kernel with only positive parts would represent a system without adaptation, while a kernel with positive and negative parts of similar area displays near perfect adaptation. In the case of WCs and CCs, both display filters with adaptation; moreover, both filters have the same shape (Fig. 3b). This suggests homology between the sensorimotor transformations of the WCs and CCs.

As in,[27] the optogenetic illumination is normalized by its mean to be between -1 and 1, and unitless. All the equations were implemented numerically in custom code written in MATLAB (https://github.com/LuisM6/ThermotaxisCrossInhibition2020). The filters were not smoothed, they were used as shown to convolve the input, this explains the high frequency components in the model results.

#### Linear combination of cooling and warming pathways

To address how the simultaneous outputs of WCs and CCs, we used the kernels obtained via optogenetics ($Hcc_{turn}, Hwc_{turn}$) to transform the unitless normalized calcium responses ($Scc_{norm}, Swc_{norm}$) into the turning rates ($Rcc_{turn}, Rwc_{turn}$) that each pathway can generate. This proceeds as follows:

$$Rcc_{turn}(t) = \int_{0}^{\infty} Hcc_{turn}(\tau) Scc_{norm}(t-\tau) d\tau \tag{13}$$

$$Rwc_{turn}(t) = \int_{0}^{\infty} Hwc_{turn}(\tau) Swc_{norm}(t-\tau) d\tau \tag{14}$$

Then, we asked whether a linear combination could capture the polarity and relative magnitude of the contribution of each sensor type to the behavioral responses to temperature fluctuations at different baselines. In other words, if we could approximate the turning rate ($R_{turn}(t)$) with the following equation:

$$R_{turn}(t) = w_{CC}(Rcc_{turn}(t)) + w_{WC}(Rwc_{turn}(t)) \tag{15}$$

The scalar weights $w_{CC}$ and $w_{WC}$ could be independent or not. We used the wildtype responses to the fast sine waves at the three baselines to fit the scalar weights via multiple linear regression. In these fitted values, when $w_{CC}$ has a high value, $w_{WC}$ has a smaller value, and vice-versa. This is to say, we observed that an increase in the CCs contribution resulted in a decrease of the WCs contribution.

The differences in the three architectures considered in the paper follow constrains that are applied to $Rcc_{turn}$ and $Rwc_{turn}$:

In the *Labeled Lines Model (LLM)*, turning during cooling can only be modulated by the CCs and turning during warming can only be modulated by the WCs. These conditions amount to a rectification of the $Rwc_{turn}$ during cooling and $Rcc_{turn}$ during warming.

In the *Cross-Inhibition Model (C-IM)*, turning is controlled by both WCs and CCs. The contributions of WCs and CCs ($Rwc_{turn}$ and $Rcc_{turn}$) to behavior are allowed to take negative values and therefore do not require a transformation to positive values nor a saturation before the linear combination. Turning rates cannot be negative though, so the result of the linear combination is transformed with a linear function into non-negative values.

In the *Cross-Activation Model (C-AM)* turning is controlled by both WCs and CCs. The contributions of WCs and CCs to behavior ($Rwc_{turn}$ and $Rcc_{turn}$) are transformed with a linear function to have non-negative values. The result of the linear combination will be always positive so it does not need to be transformed.

In all cases, the values of the weights ($w_{CC}$ and $w_{WC}$) are linearly regressed to match the amplitude of the wildtype animal responses. See the Extended Methods for a detailed derivation of the models.

In the case of the *C-IM*, to capture the relationship between the relative contributions of both types of sensors, we reparametrized the scalar weights as follows:

$$w_{CC} = k(w) \tag{16}$$

$$w_{WC} = k(1-w) \tag{17}$$

With this parametrization $w$ captures the relative contribution of each sensor type. Here, $w$ is set to vary from 0 to 1, with 0 being the case where $w_{CC} = 0$ and therefore the contribution of the CCs is 0 and behavior is preferentially driven by the WCs. As $w$ increases, the contribution of the CCs increases and the contribution of the WCs decreases. When $w$ is 0.5, $w_{CC} = w_{WC}$ and the contribution of WCs and CCs is the same. When $w$ is 1, $w_{WC} = 0$ and therefore the contribution of the WCs is 0 and behavior is preferentially driven by the CCs. The scalar parameter $k$ is fixed and sets the scale of the predicted turning responses.

This parametrization allows to use two successive linear regressions instead of a multiple linear regression. In the first linear regression, we fit the $w$ parameter at different temperature baselines normalizing both the predicted and the experimentally measured responses by their peak response. In the second linear regression, we fit $k$ to match the amplitude of the experimentally measured responses. While $w$ is fitted for each temperature baseline, $k$ is a fixed parameter, fitted for all conditions at the same value.

These parametrizations are fitted to data using the MATLAB Curve Fitting Toolbox.

**Incorporating the baseline turning rate**

We consider the turning rate in the absence of WCs and CCs (obtained experimentally with the Ir93aMI mutants) the baseline turning rate ($R_{bln}(t)$). The baseline turning rate is incorporated to the turning rate produced by the WCs and CCs pathway as an additive correction:

$$R_{turn}(t) = w_{CC}(Rcc_{turn}(t)) + w_{WC}(Rwc_{turn}(t)) + R_{bln}(t) \tag{18}$$

### Optogenetics at different temperatures

The efficiency of CsChirmson to induce neural activity has not been tested as a function of temperature. In our attempt to test the behaviors produced by CCs and WCs we found that at higher temperatures CsChrimson is less effective. For example, labeling Or49a (a wasp pheromone sensing neuron[59]) we observed that the turning rate is drastically diminished at higher temperatures using an identical light intensity ramps (Supplementary Fig. 9A). This effect can be alleviated by increasing the light intensities used in the experiment (Supplementary Fig. 9B). Therefore, we had to use higher light intensities at high temperatures to test whether CCs and WCs are capable of modulating the turning rate in that range.
