## Supplementary Material Part 1 for "Synchronous and opponent thermosensors use flexible cross-inhibition to orchestrate thermal homeostasis"

### Ir93a and Ir25a are expressed in the WCs

In a previous study, it was shown that Ir93a is expressed in 5 neurons in each DOG including the 3 CCs.[18] We asked whether the other two neurons are the WCs. We immunostained the DOG with anti-Ir93a in an *Ir68a-Gal4;UAS-GFP* larva, thus confirming that the WCs express Ir93a (Supplementary Fig. 2D). A previous study established complete overlap of Ir25a and Ir93a expression in the DOG using immunostaining.[48] Thus, Ir25a is also expressed in the WCs.

### WC and CC morphologies

The morphology of the larval WCs has homologies to warming cells identified in the arista of the adult.[19] One difference is that the larval WCs have an additional thin dendritic process that projects to the larval surface through a pore in the olfactory dome (Supplementary Fig. 4A, B). This dendritic process is much thinner than that of olfactory neurons that also project to the olfactory dome. This can be observed clearly from the EM cross-sections at the base of the dome (Supplementary Fig. 4C).

The A- and B-CCs have differences in the structures of their outer segments. The A-CCs have a larger bulb (Supplementary Fig. 4A). This feature might contribute to the difference in thermosensitivity between A- and B-CCs. Previous studies have identified correlations between the extent of lamellation in the thermosensory neurons of different insects and their thermosensitivity.[54]

The cell body and outer segment of the B-CC runs parallel to an unknown non-thermosensory cell in the DOG. This cell's outer segment is morphologically different than that of the WCs (Supplementary Fig. 4A). The unknown cell also sends a thin dendritic process to the olfactory dome pores (Supplementary Fig. 4B). We were unable to find thermosensory responses in any other cells but the 2 WCs and 3 CCs in the DOG using pebbled-Gal4/UAS-GCaMP6m to label neurons (Supplementary Fig. 1). We also were unable to find potential Ir21a, Ir68a, or Ir93a expression in any cell except the 3 CCs and 2 WCs.

### Ectopic expression of cooling and warming receptors in opponent cells

We used UAS-Gal4 to ectopically express the cooling receptor Ir21a in the warming cells (UAS-GCaMP6m/UAS-Ir21a; Ir68a-Gal4/+), converting the WCs into cooling cells (Supplementary Fig. 5A). We also ectopically expressed Ir68a in the cooling cells (11F02-Gal4/+; UAS-GCaMP6m/UAS-Ir68a), and found that the cooling responses of the CCs were drastically diminished (Supplementary Fig. 5B, C). The differences in these physiological phenotypes may be caused by differences in receptor competition in these cell types.

### The emergence of cross-inhibition

In our model, WCs and CCs both activate and cross-inhibit the turning responses to cooling and warming. The relative weights of activation and cross-inhibition determine whether the larva avoids cooling at low temperatures or avoids warming at high temperatures. These avoidance responses drive movement towards the homeostatic set-point, where the relative weights of activation and cross-inhibition lead to cancellation of turning responses.

Cross-inhibition emerges from the polarity of the sensors themselves, their bidirectional phasic responses, and the linear model that transforms WC and CC activity into turning responses. Here, we describe how this works. First, consider the WCs. Cooling inhibits the WCs at all temperatures (Fig. 3D-F). Moreover, activation and inhibition of the WCs is associated with higher and lower turning rates at all temperatures (Fig. 5A, B). When the larva experiences cooling at low temperatures, WCs are inhibited (Fig. 3D). This lowers the turning response to cooling at low temperatures (Fig. 6B). But CCs are also activated by cooling at low temperatures. Because the overall turning response is a weighted sum of WC and CC activities, cooling avoidance emerges because the turning response to cooling caused by the CCs outweights the cross-inhibition of the turning response to cooling caused by the WCs (Fig. 4H). Similar patterns explain the response to cooling and warming at all temperatures (Fig. 6B-D). The only free variable in the model are the relative weights of CC and WC output in determining turning responses.

The flexibility or change in the scalar weights ($w_{WC}$ and $w_{CC}$) of the cross-inhibition is asymmetric. At low temperatures $w_{WC}$ is 2x smaller than $w_{CC}$, and at high temperatures $w_{WC}$ is 4x larger than $w_{CC}$ (Fig. 4G). This asymmetry is caused by reduced sensitivity of the WCs at high temperatures (Fig. 3F) compared to the A-CCs. However, regardless of the loss in sensitivity, the WCs still make a statistically significant contribution to warming avoidance to fast stimulus changes at high temperatures (Fig. 6D).

### Correlation between model predictions and experiments

The model captures a large percentage of the behavioral response dynamics. Most of the responses to slow and fast sine waves of temperature in the three thermosensory contexts results in correlation coefficients between 0.65 and 0.95 for wildtype and mutant larvae (Fig 6E, and Supplementary Fig. 9D).

### Turning rate additional features

Each turn event has different properties like the number of head sweeps it involves and their size or duration. These variables are less sensitive to temperature changes (Supplementary Fig. 12). However, in the cases where wildtype and mutant larvae display statistically significant differences, they support the turning rate tendency. For example, the turn size at low temperatures is higher for larvae defective for WCs function, consistent with the mutants being more sensitive to cooling because the WCs inhibition is absent (Supplementary Fig. 12A). Wildtype animals make more head sweeps than WC-defective mutants at high temperatures (Supplementary Fig. 12B), consistent with WCs contributing to warming avoidance at high temperatures. Wildtype animals perform longer turns during cooling than CC-defective mutants at low and ideal temperatures (Supplementary Fig. 12A), consistent with the role of the CCs in cooling avoidance. Mutants defective for both WCs and CCs have shorter cooling induced turns at ideal temperatures and also make fewer head sweeps (Supplementary Fig. 12A-C), consistent with the WCs and CCs being the main drivers to thermosensation in the larval head.

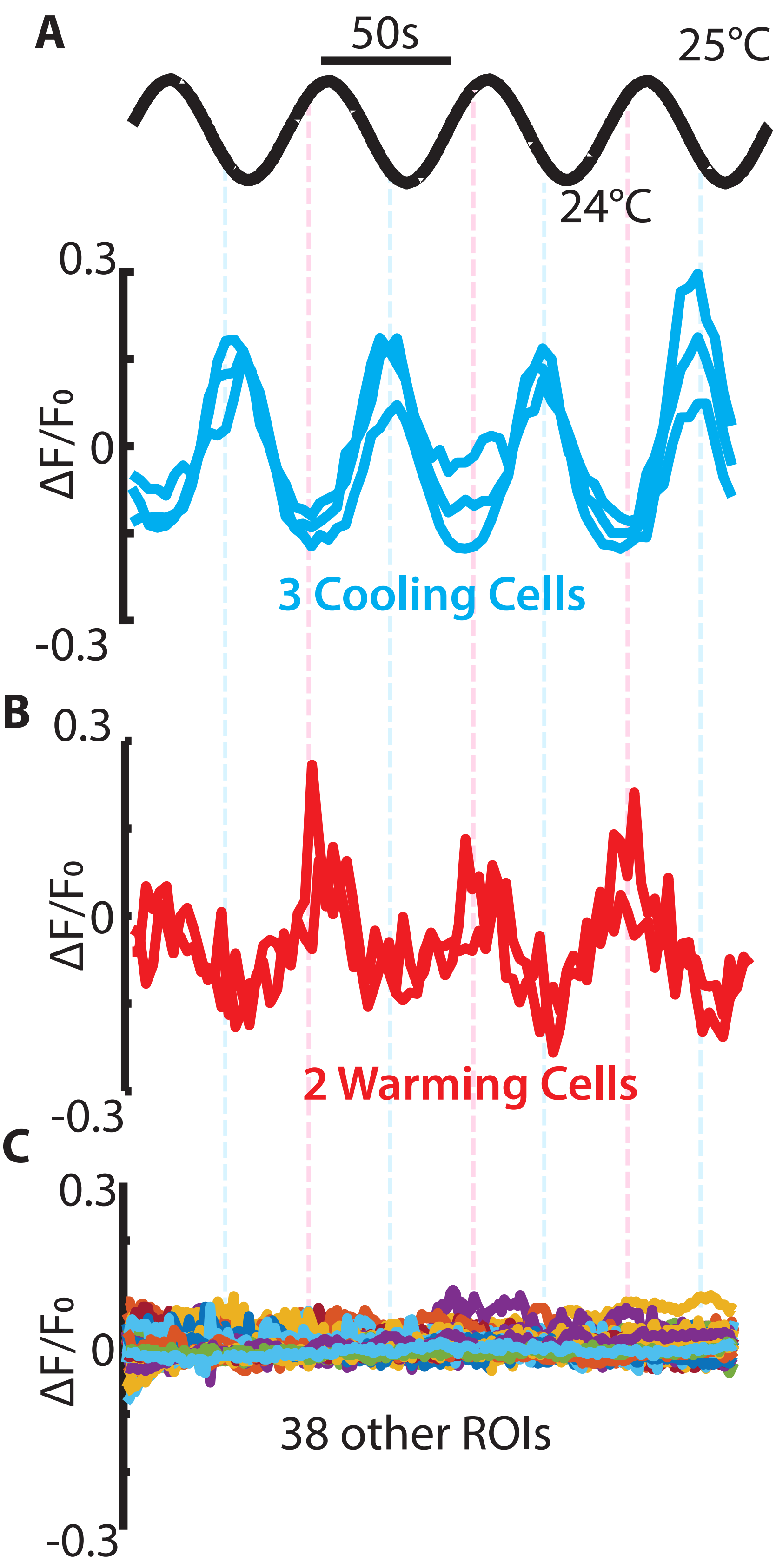


**Supplementary Figure 1. | Thermosensors in the larva head. A**, Responses recovered in the Dorsal Organ Ganglion (DOG) via CNMF of the CCs (in cyan), (**B**) the new WCs (in red), and (**C**) the other 38 ROIs that were not responsive to temperature. (Genoype: $w^{1118}$;Pebbled-Gal4/UAS-GCaMP6m, n=6).

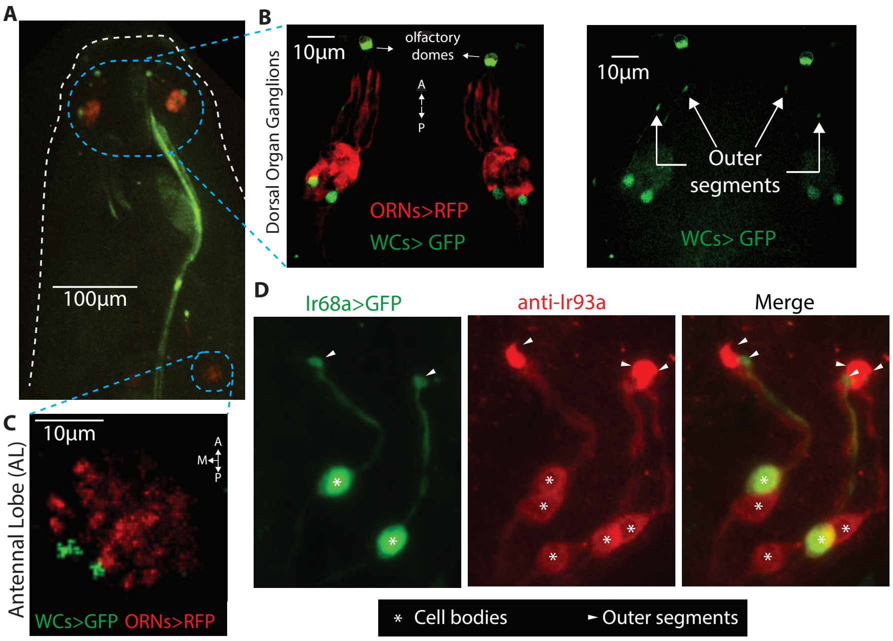


**Supplementary Figure 2.** | **Ir expression in the Dorsal Organ Ganglion. A**, Ir68a-Gal4 expression pattern in the larval body. Ir68a-Gal4 drives UAS-CD8-GFP expression and the orco drives RFP expression in the olfactory receptor neurons(ORNs). (Genoype: UAS-CD8-GFP;orco-RFP/Ir68a-Gal4). **B**, Location of the WCs cell bodies relative to the ORNs. The olfactory domes are autofluorescent. Removing the red channel, we can better observe the outer segments of the WCs (right inset). **C**, The axon terminals of the WCs form two separate glomeruli in the antennal lobe and are located medial, dorsal and posterior relative to the ORNs glomeruli. **D**, Immunostaining with anti-Ir93a shows expression in 5 cells, two of them overlap with the WCs. The location of the outer segments of WCs relative to the other Ir93a expressing neurons can be observed in the merged image.

A

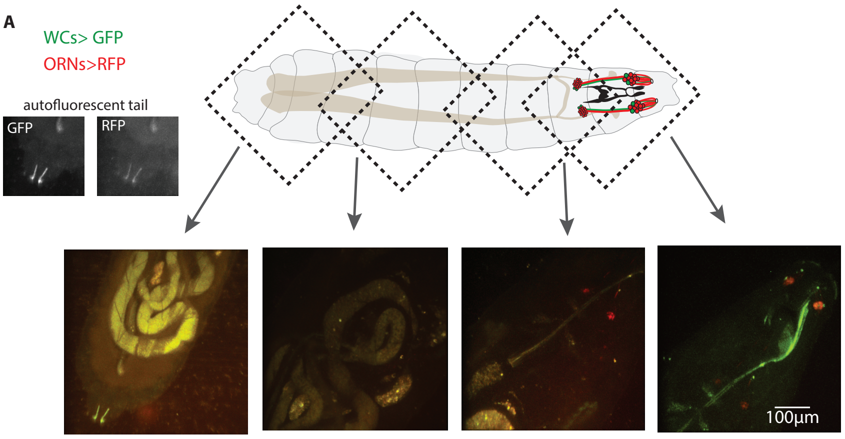


B

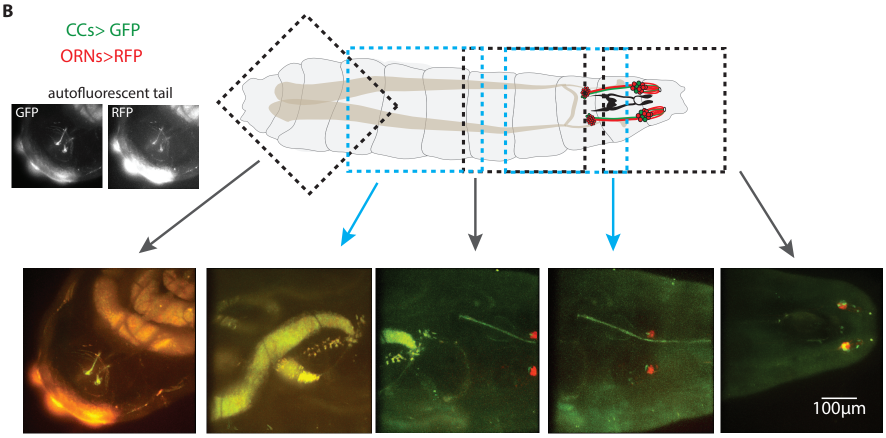


**Supplementary Figure 3.** | **Gal4 driver expression patterns. A**, Expression pattern of Ir68a>Gal4 in the entire larva. (Genoype: UAS-CD8-GFP;orco-RFP/Ir68a-Gal4). Green fluorescence is only detected in the WCs in the DOG. No other neurons present green fluorescence. Some structures like the intestines and the tail are autofluorescent (inset) **B**, Expression pattern of R11F02>Gal4 in the entire larva. (Genoype: UAS-CD8-GFP/R11F02-Gal4;orco-RFP/+). Green fluorescence is only detected in the CCs in the DOG. No other neurons present green fluorescence. Some structures like the intestines and the tail are autofluorescent (inset)

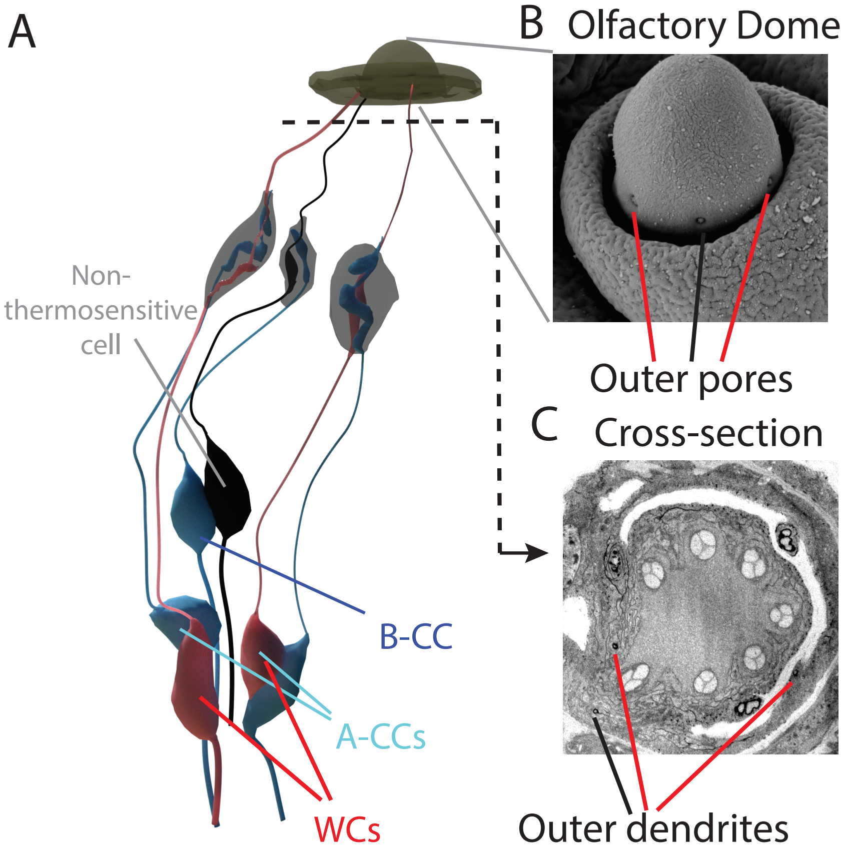


**Supplementary Figure 4.** | **Morphology of the thermosensory cells. A**, Electron microscopy reconstruction of the WCs and CCs cell bodies and outer segments. **B**, Detailed view of the olfactory dome with the three pores, two of them innervated by a dendritic process of the WCs. **C**, Cross-section at the base of the olfactory dome.

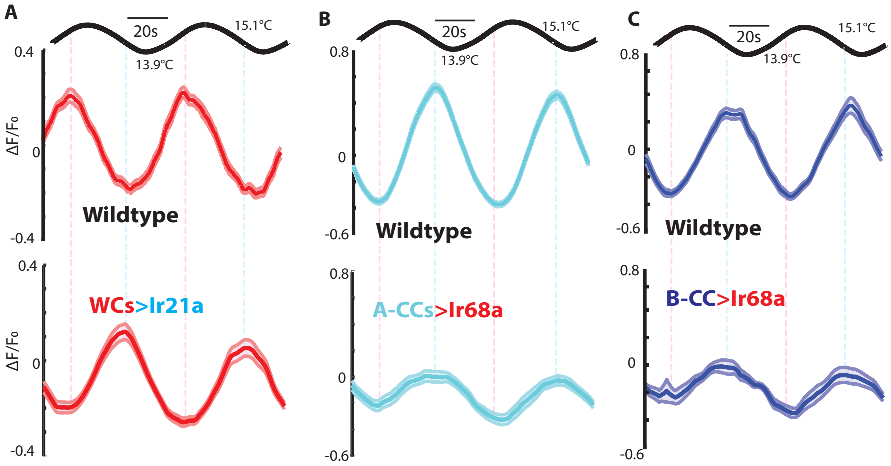


**Supplementary Figure 5. | Ectopic expression of opposed receptors. A**, Ectopic expression of Ir21a in the WCs inverts their polarity by transforming them into cooling activated sensors. (Genoype: $w^{1118}$;UAS-Ir21a;Ir68a-Gal4, UAS-GCaMP6m, n=8). **B**, **C**, Ectopic expression of Ir68a in the CCs attenuates their cooling response. (Genoype: $w^{1118}$;R11F02-Gal4;UAS-Ir68a/UAs-GCaMP6m, n=6). Shaded regions are the standard error of the mean (s.e.m.).

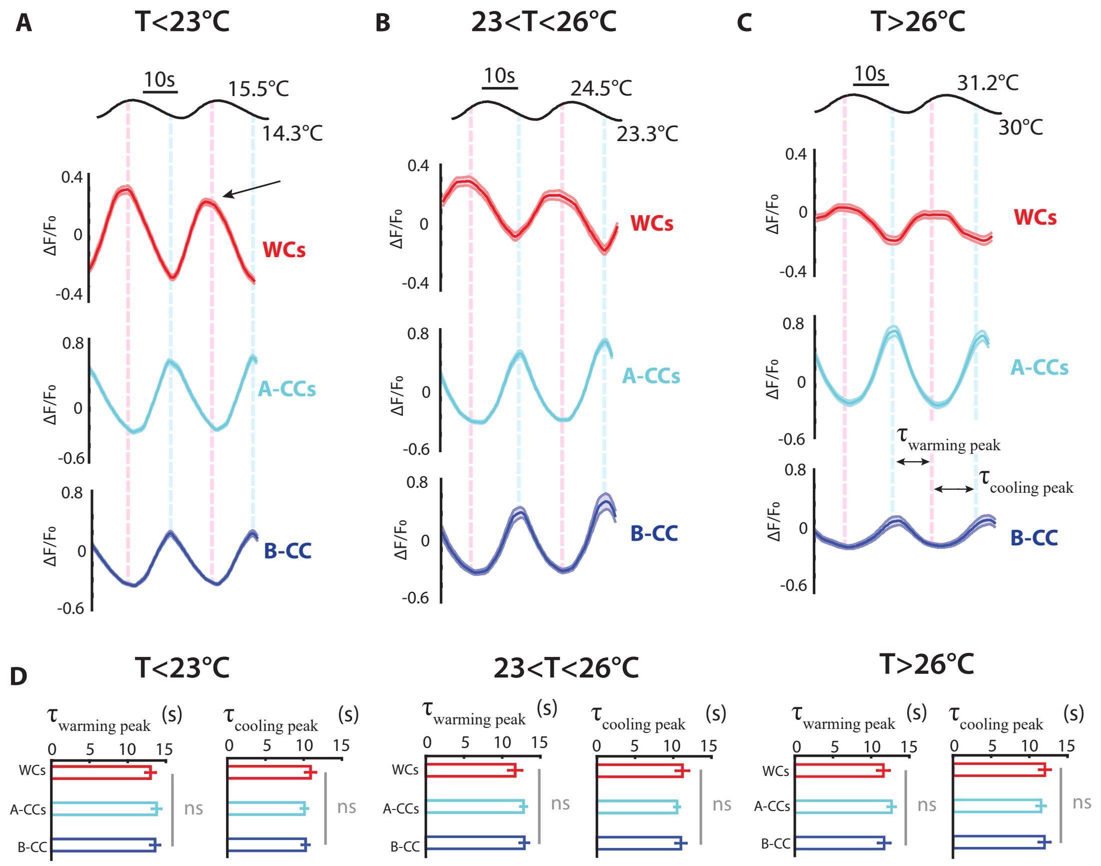


**Supplementary Figure 6.** | **WCs and CCs are synchronous and opponent thermosensors.A**, **B**, **C** *Drosophila* larvae expressing GCaMP6m in the WCs (UAS-GCaMP6m;Ir68a-Gal4) or CCs (R11F02-Gal4; UAS-GCaMP6m) calcium responses to temperature sine waves of 1.2°C amplitude. This is the same dataset shown in Fig. 3. **D** The peak times during cooling or warming of WCs and CCs responses are not significantly different (Kruskal-Wallis test, error bars are the s.e.m) and their polarities are always opposed. WCs are activated by warming and inhibited by cooling. CCs are inhibited by warming and activated by cooling.
