## Supplementary Material Part 2 for "Synchronous and opponent thermosensors use flexible cross-inhibition to orchestrate thermal homeostasis"

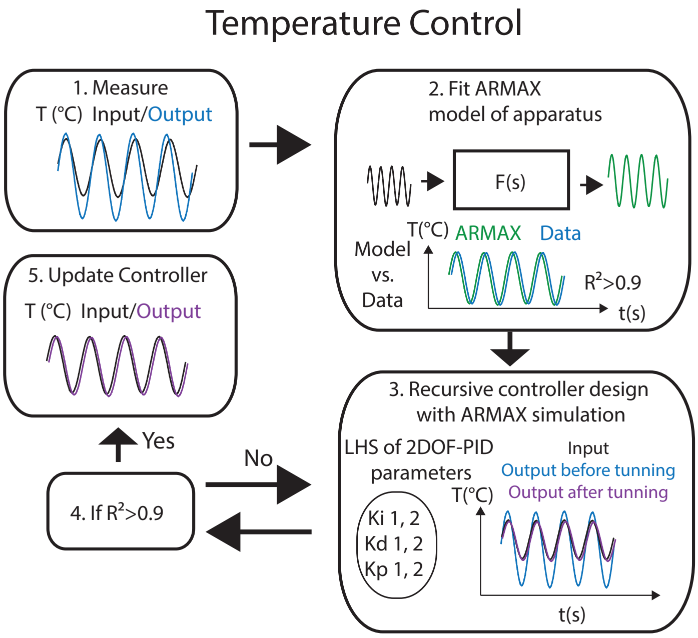


**Supplementary Figure 7.** | **Temperature control technique.** Algorithm to calibrate the temperature controllers using an ARMAX systems identification technique. First we measure the input-output relationship of the device. Then, we use that data to fit an ARMAX model to the device functioning. Next, we scan parameter combinations of the second degree of freedom of the PID control and analyze their performance using the ARMAX model. Once a parameter combination produces convergent results, that parameter combination is selected and tested experimentally.

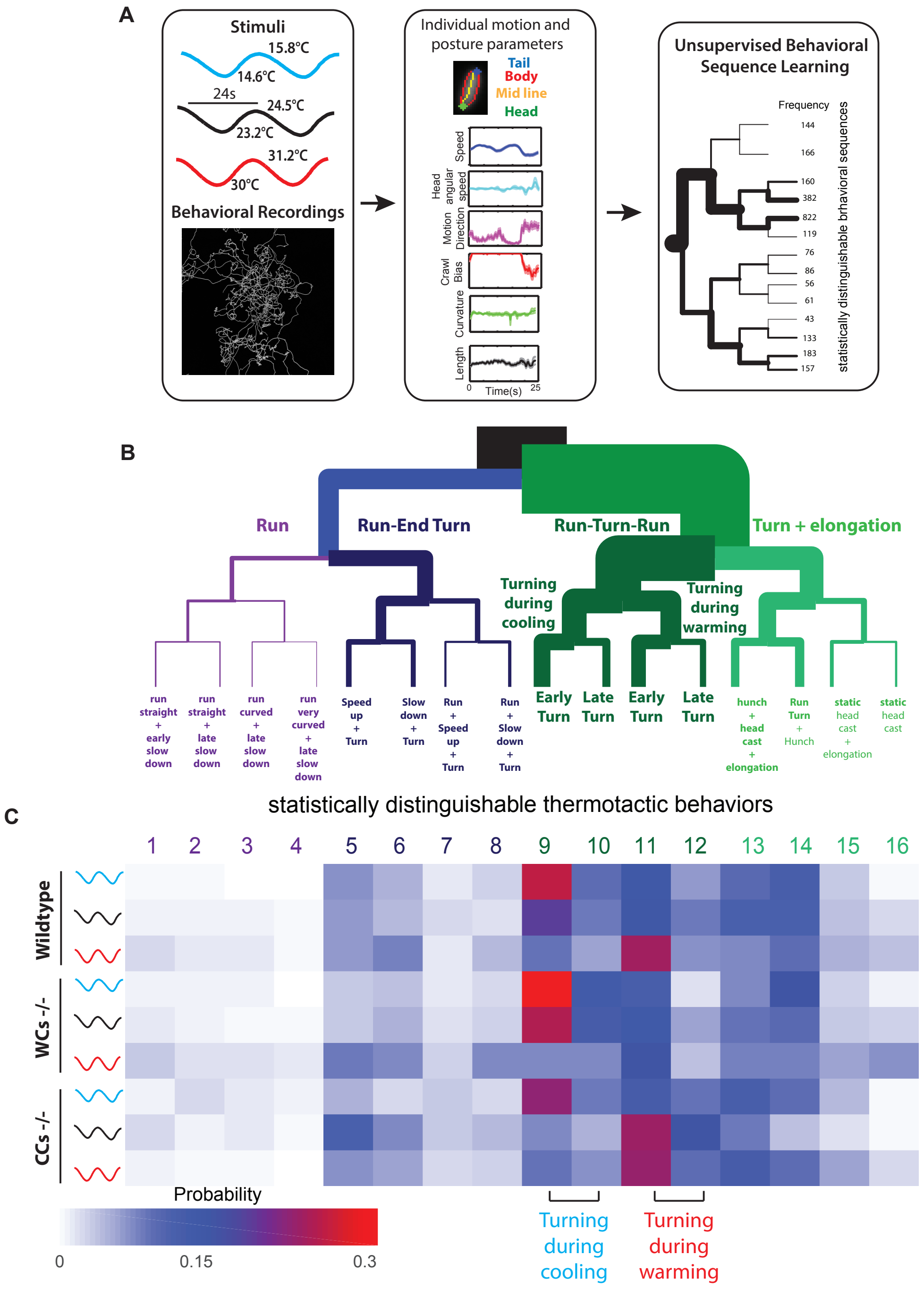


**Supplementary Figure 8.** | **Unsupervised behavioral sequence learning. A**, Behavioral analysis pipeline. First we collected movies of freely crawling larvae exposed to sinusoidal stimuli for 20 minutes. Then, we segmented these movies to identify individual larvae and their posture. We calculated relevant behavioral features and finally we use these time traces as the input for the iterative denoising trees behavioral classifier. **B**, The resulting behavioral tree for wildtype and mutant larvae exposed to sine waves at three different baselines. The labels of each behavior are assigned post hoc based on the time series of behavioral features in each branch. **C**, Heatmap of probabilities of displaying each statistically distinguishable behavioral sequence. Only behaviors 9 to 12 come from different distributions at different baselines using Steel-Dwass test with $p<0.01$.

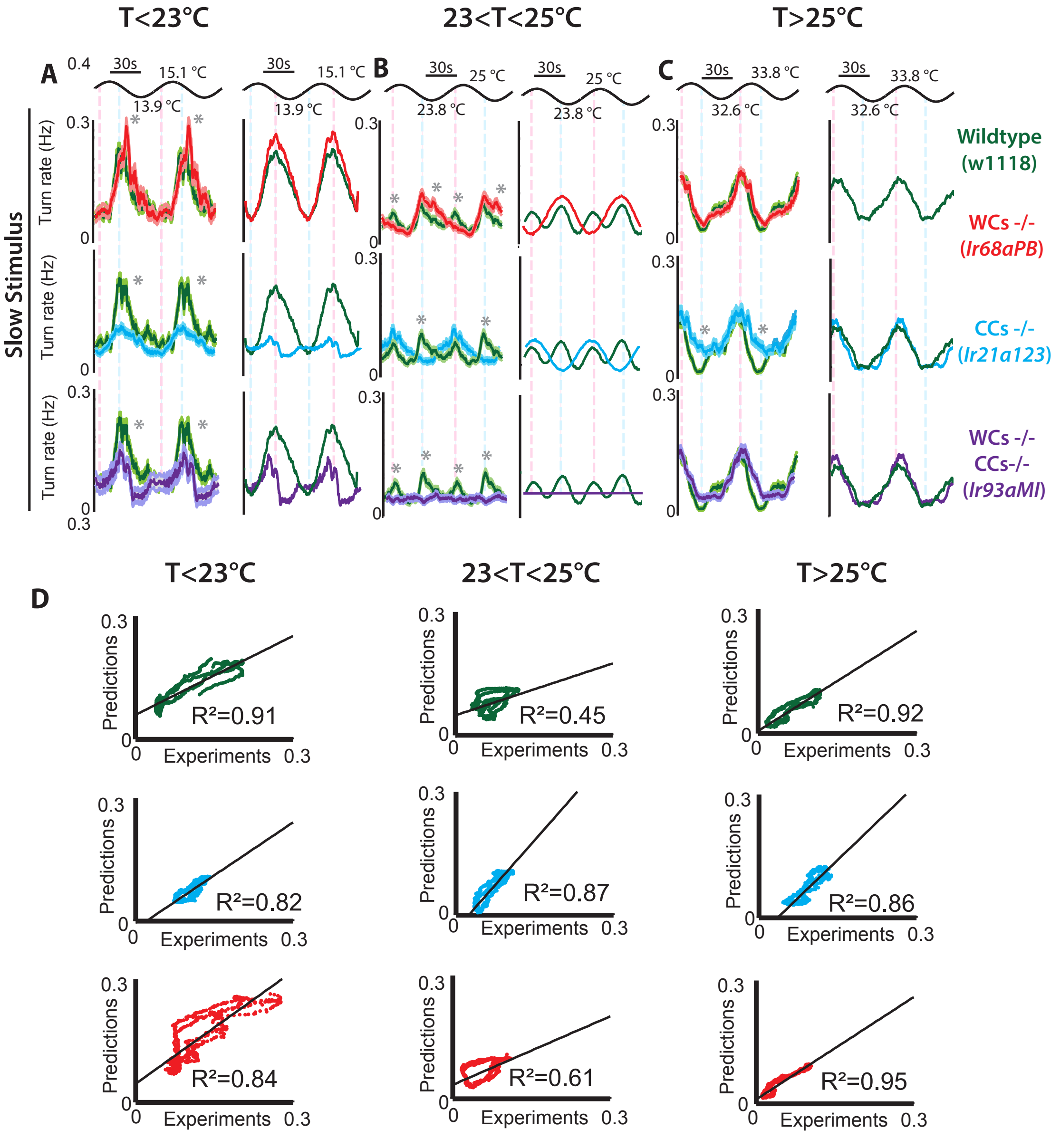


**Supplementary Figure 9.** | **Behavior dataset for slow temperature fluctuations. A-C left**, Experimental results of the turning rate response to slow sinusoidal waves of temperature of wildtype larvae ($w^{1118}$ in green), larvae defective for CCs' function ($w^{1118}$; $Ir21a^{123}$ in cyan, larvae defective for WCs' function ($w^{1118}$; $Ir68a^{PB}$ in red), and larvae defective for WCs and CCs function ($w^{1118}$; $Ir93a^{MI05555}$ in purple at ambient temperatures below (**A**), near (**B**), and above (**C**) the homeostatic set-point. Shaded regions are the s.e.m. * indicate different mean turning rate during cooling or warming using Chi-squared test with Bonferroni correction ($p<0.005$). **A-C right**, Quantitative predictions of the turning rate response to a slow sinusoidal wave of temperature . Wildtype larvae's turning rate in green, larvae defective for WCs' function in red, larvae defective for CCs' function in cyan, and larvae defective for WCs and CCs function in purple at ambient temperatures below (**A**), near (**B**), and above (**C**) the homeostatic set-point. $w^{1118}$ in green, n=59 in **A**, n=75 in **B**, and n=72 in **C**. $w^{1118}$; $Ir21a^{123}$ in cyan, n=70 in **A**, n=73 in **B**), and n=81 in **C**. $w^{1118}$; $Ir68a^{PB}$ in red, n=78 in **A**, n=73 in **B**, and n=77 in **h**. $w^{1118}$; $Ir93a^{MI05555}$ in purple, n=65 in **A**, n=59 in **B**, and n=70 in **C**. **D**, Correlation plots corresponding to the curves in **A-C**. The first column corresponds to the sinusoidal waves at temperatures below the set-point, the second one to the sinusoidal waves near the set-point, and the third one to the sinusoidal waves above the set-point.

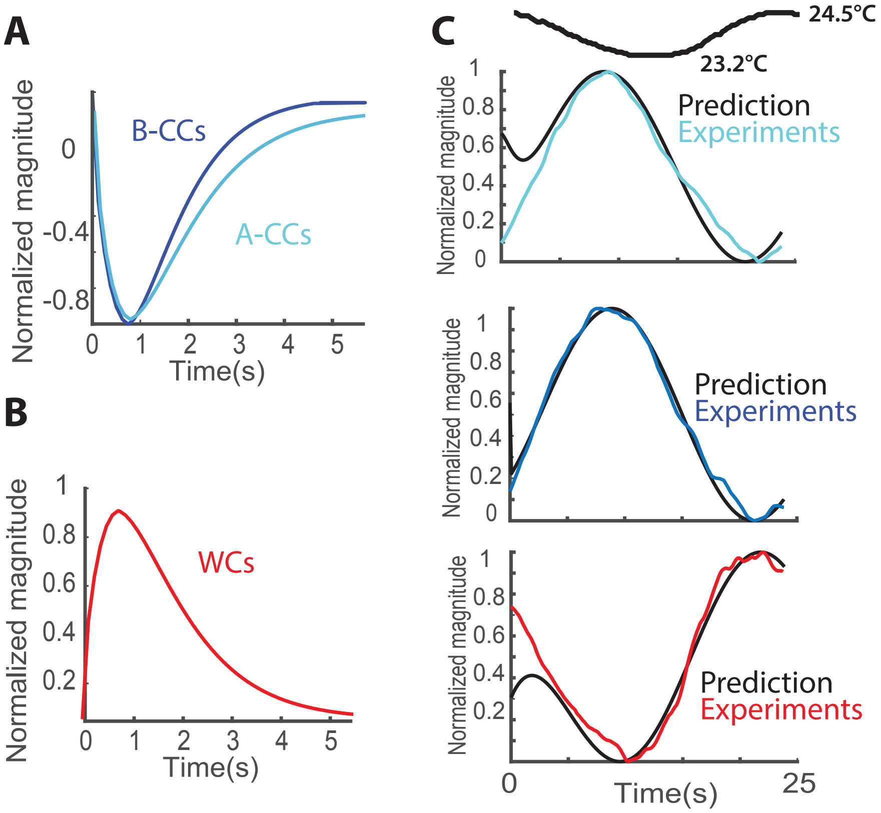


**Supplementary Figure 10.** | **Neural and behavioral linear filters. A**, Linear filters fitted to the sine wave calcium responses of Fig. 2. The three filters estimate well the shape of the responses and show that the A and B-type CCs operate in the same timescale. **B**, We calculate the transfer function from optogenetic stimulus to turning rate. The linear filters of the WCs (in red) and CCs (in cyan) have the same shape. (Genotypes: UAS-CsChrimson;Ir68a-Gal4 n=122 , 11F02-Gal4; UAS-CsChrimson n=117 )

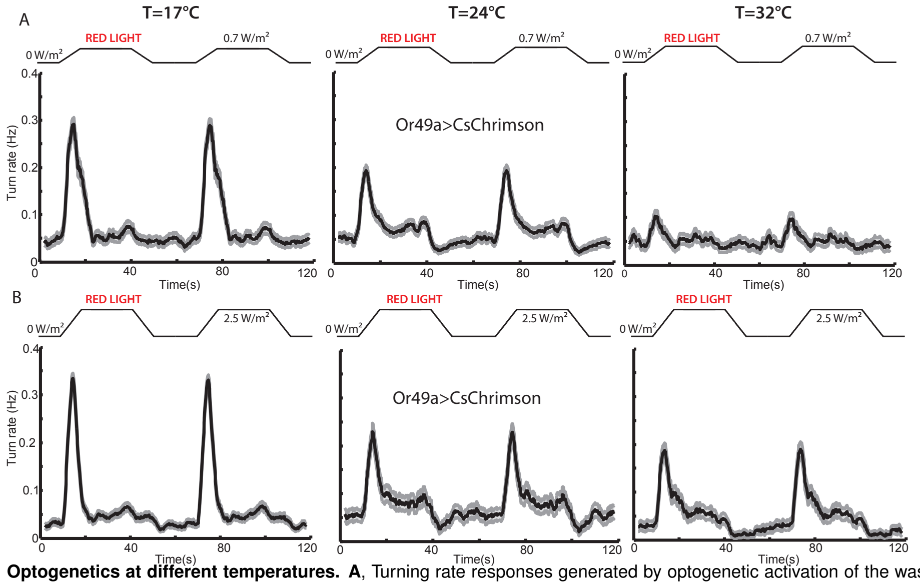


**Supplementary Figure 11.** | **Optogenetics at different temperatures.** **A**, Turning rate responses generated by optogenetic activation of the wasp pheromone sensor Or49a, using 0.7W/m$^2$ of red light intensity. **B**, Turning rate responses generated by optogenetic activation of the wasp pheromone sensor Or49a, using 2.5W/m$^2$ of red light intensity.(Genoype: w$^{1118}$;Or49a-Gal4/UAS-CsChrimson, n=90-118 per temperature and light intensity).

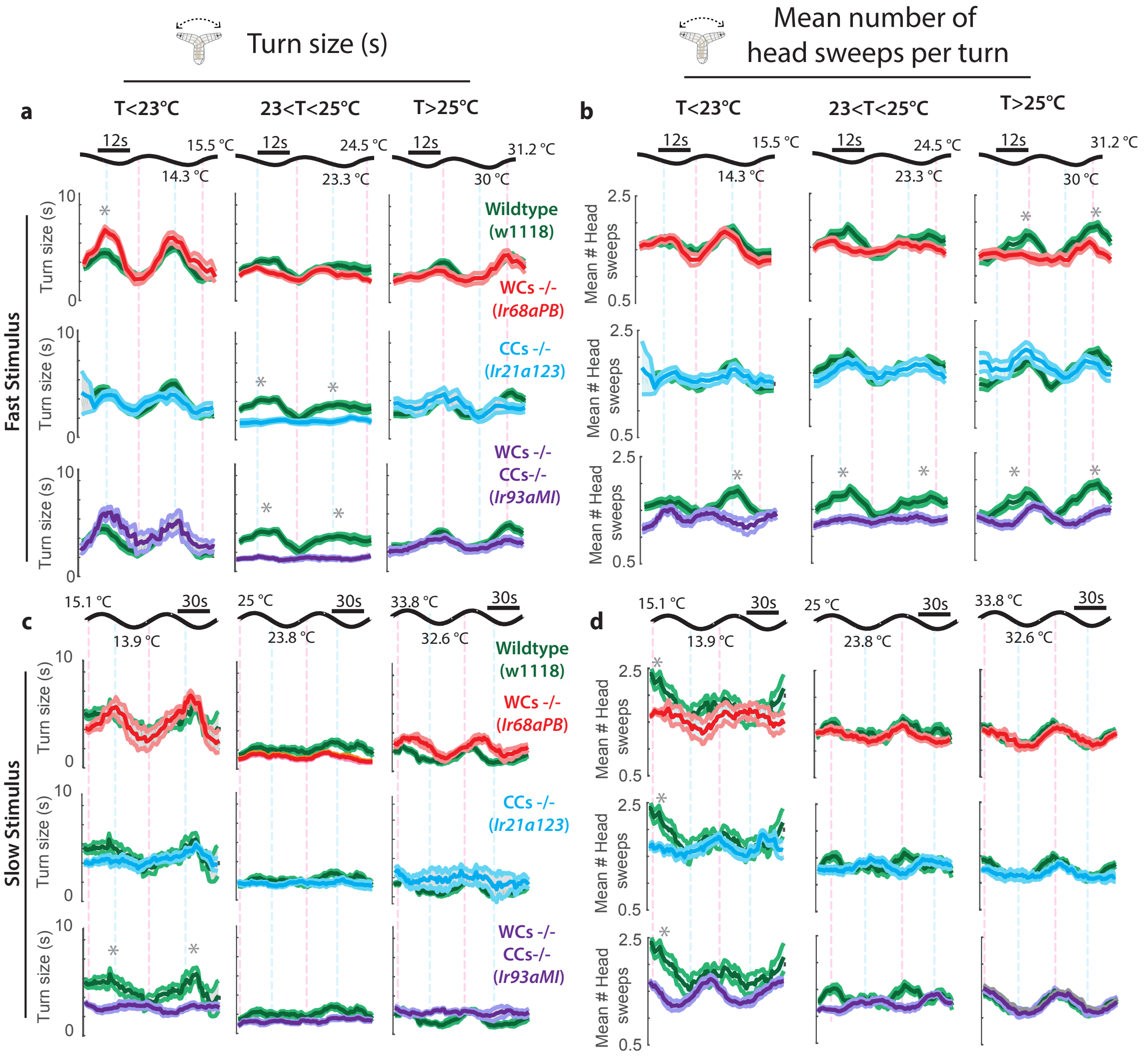


**Supplementary Figure 12.** | **Properties of the turn events.** **A**, **C**. Size of each turn in seconds for fast (**A**) and slow (**C**) temperature sine waves, below (first column), near (second column), or above (third column) preferred temperatures. Mutants defective for the WCs are in red, mutants defective for the CCs in cyan, mutants defective for both WCs and CCs in purple, and wildtype animals in green. the number of animals and genotypes match the ones in Fig. 4. The shaded regions around the curves is the s.e.m. * indicate different mean turning size during cooling or warming using Steel-Dwass test with p<0.01. **B**, **D**. Number of head sweeps in each turn for (**B**) and slow (**D**) temperature sine waves, below (first column), near (second column), or above (third column) preferred temperatures. Mutants defective for the WCs are in red, mutants defective for the CCs in cyan, mutants defective for both WCs and CCs in purple, and wildtype animals in green. the number of animals and genotypes match the ones in Fig. 4. The shaded regions around the curves is the s.e.m. * indicate different mean turning size during cooling or warming using Steel-Dwass test with p<0.01.
